# Structural features and conservation of the lncRNA HOTAIR in primates

**DOI:** 10.64898/2026.05.18.725451

**Authors:** Zion R. Perry, Anish Beeram, Logan Lee, Anna Marie Pyle

## Abstract

Long noncoding RNAs (lncRNAs) regulate diverse cellular processes and are frequently implicated in disease, but their functional mechanisms often remain elusive. One such lncRNA, HOTAIR (Hox transcript antisense intergenic RNA), is a ∼2.1 kb mammalian transcript whose overexpression promotes invasion and metastasis in breast cancer. However, the mechanisms by which HOTAIR influences gene regulation in cancer are poorly understood. To approach this problem through a structural lens, we predicted the full-length *in cellulo* secondary structure of HOTAIR using chemical probing in a metastatic breast cancer cell line. The resulting structure shows that HOTAIR adopts a multidomain architecture and has local structural features unique to the cellular context. Comparison between *in vitro* and *in cellulo* chemical probing identifies regions of differential accessibility that may indicate context-dependent molecular interactions or folding. Conservation analyses further reveal that HOTAIR is conserved across primates with evidence of structural covariation in specific domains. Together, these results provide a roadmap for future mechanistic studies of structure-function relationships in HOTAIR and its contribution to gene regulation in cancer.

## INTRODUCTION

The classical view of RNA as a messenger for protein translation represents only part of RNA’s complex roles in cells. Noncoding RNAs (ncRNAs) participate in many essential cellular processes, and their mis-regulation is associated with numerous human diseases. Long noncoding RNAs (lncRNAs) are a class of ncRNAs longer than 500 nucleotides that are typically transcribed by RNA polymerase II and often capped, spliced, and polyadenylated (Mattick et al. 2023). Since the discovery of the first human lncRNA Xist in 1992 (Brown et al. 1992), studies of lncRNAs have revealed their diverse regulatory mechanisms at all stages of gene expression. LncRNAs may function *in cis* at their locus of transcription, such as through transcriptional interference or enhancer-like activity, or function *in trans* at distant chromatin or cellular locations (Rinn 2014; Kopp and Mendell 2018). In some cases, the underlying DNA locus may function in transcriptional regulation (Engreitz et al. 2016) or the lncRNA may encode for short peptides or proteins (Kawashima et al. 2003; Li et al. 2020). Despite rapid advances in the lncRNA field, the functional mechanisms of most lncRNAs still remain poorly understood.

The lncRNA HOX transcript antisense intergenic RNA (HOTAIR) is a prominent example for which we have an incomplete mechanistic understanding. HOTAIR was first discovered by through microarray sequencing of the *HOX* loci in human fibroblasts, which revealed a transcript antisense to the HOXC locus on chromosome 12 that regulates gene expression *in trans* at the HOXD locus on chromosome 2 (Rinn et al. 2007). Knockdown of HOTAIR reduced histone 3 lysine 27 trimethylation (H3K27me3) and decreased occupancy of the Polycomb Repressive Complex 2 (PRC2) subunit Suz12 at HOXD promoters, implicating HOTAIR in epigenetic silencing. Subsequent studies identified the binding sites for PRC2 near the 5’ end of HOTAIR and for another chromatin silencing complex, Lysine-Specific Demethylase 1 (LSD1), near the 3’ end (Tsai et al. 2010). This led to the model of HOTAIR as a molecular scaffold for coordinating chromatin-modifying complexes. However, later studies found that HOTAIR can mediate transcriptional repression independently of PRC2 binding (Portoso et al. 2017), suggesting that recruitment of chromatin modifiers may occur indirectly and after HOTAIR localization to target genes (Brockdorff 2013). Additional studies have since reported interactions between HOTAIR and diverse protein partners (Meredith et al. 2016; Delhaye et al. 2022; Porman et al. 2022), indicating HOTAIR may have multifunctional roles in gene expression. Although HOTAIR participates in epigenetic silencing at some level, the mechanisms of protein recruitment and target gene regulation remain unresolved.

Although HOTAIR plays a normal role in early development (Rinn et al. 2007), its aberrant overexpression has been extensively linked with cancer progression. In breast cancer, HOTAIR is highly overexpressed in metastatic tumors relative to primary tumors and normal tissue (Gupta et al. 2010). Overexpression of HOTAIR in breast cancer cell lines confirmed its invasive influence in cell culture and mouse models, notably in MDA-MB-231 cells, a triple negative and highly invasive breast cancer cell line. Other lncRNAs also exhibit gain of function, oncogenic phenotypes in various cancer pathways, and HOTAIR serves as one such oncogenic lncRNA for dissecting how RNA-mediated regulatory mechanisms drive tumor progression (Mozdarani et al. 2020; Winkler and Dimitrova 2022). HOTAIR’s oncogenic effects are associated with many cancers, including liver (Yang et al. 2011), colorectal (Kogo et al. 2011), lung (Loewen et al. 2014), and ovarian (Dong and Hui 2016) cancers. Breast cancer thus serves as a model system to study paradigms of lncRNA dysregulation that may have broader application.

RNA structure is central for regulating its function. Although there are no high-resolution tertiary structures of any lncRNA, secondary structure is a tractable starting point for discovering lncRNA structural modules that may aid functional studies or 3D structure determination of RNA-protein complexes. An initial indication of structure-function relationships in HOTAIR came from an 89-nucleotide minimal PRC2-binding domain (212-300 nt), where nuclease mapping experiments were used to constrain a secondary structure prediction of two stem structures proposed to be important for binding (Wu et al. 2013).

Chemical probing is another method of determining the secondary structure of RNA. Selective 2’-hydroxyl acylation analyzed by primer extension (SHAPE) reagents modify the 2’OH of the ribose sugar in the backbone in a manner proportional to its flexibility, and generally base-paired nucleotides will be less modified than single-stranded nucleotides (Siegfried et al. 2014; Smola et al. 2015b). This generates a reactivity profile that can be used to constrain *in silico* secondary structure prediction algorithms (Reuter and Mathews 2010). The first complete HOTAIR secondary structure was obtained on a full-length HOTAIR transcript that was transcribed, natively purified *in vitro*, and then probed using SHAPE with capillary electrophoresis (Somarowthu et al. 2015). This work suggested that HOTAIR folds into four modular domains, which were proposed to provide platforms for protein interactors such as PRC2 and LSD1. A more recent study also used *in vitro* transcribed HOTAIR combined with SHAPE probing to measure areas of protection upon RNA target binding (Balas et al. 2021).

Since then, advances in chemical probing and sequencing techniques have enabled more accurate RNA structure determination by combining chemical probing with mutational profiling and next generation sequencing. This methodology has been further extended to cellular contexts, and improved probes have been developed (Smola and Weeks 2018). Despite these advances, there are few published *in cellulo* secondary structures of lncRNAs except for Xist, MEG3, MALAT1, NORAD, COOLAIR, SChLAP1, and Pnky (Smola et al. 2016; Sherpa et al. 2018; McCown et al. 2019; Chorostecki et al. 2021; Yang et al. 2022; Monroy-Eklund et al. 2023; Falese et al. 2025; Oh et al. 2025; Saha et al. 2026). Although *in vitro* structures derived from purified and refolded RNA are informative, the *in cellulo* structure is more informative because molecular interactions, chemical modifications, molecular crowding, or other cellular conditions may influence folding. Particularly when carried out in parallel with *in vitro* probing on natively folded transcripts, comparative methods can yield important insights into lncRNA structure and function (Smola et al. 2015a).

LncRNAs generally exhibit more rapid sequence divergence than protein-coding genes, though they may retain conserved structures (Ulitsky and Bartel 2013; Johnsson et al. 2014; Szczesniak et al. 2021; Mattick et al. 2023). Structural conservation of lncRNAs can be measured through statistical covariation tests on multiple sequence alignments, where compensatory sequence mutations across evolution preserve base pairing (Tavares et al. 2019). Despite this, any form of structural conservation in lncRNAs has been questioned (Rivas et al. 2017; Rivas 2021). However, if one evaluates conservation and covariation within a narrower evolutionary timespan, such as the primate lineage, lncRNAs show evidence of sequence and structural conservation comparable to that of highly conserved regulatory genes (Beeram et al. 2026). Measuring lncRNA conservation requires accurate annotation across sufficiently diverse genomes, which recent primate genome assemblies now make more tractable (Kuderna et al. 2023; Kuderna et al. 2024).

In this study, we present the first, full-length *in cellulo* SHAPE-MaP secondary structure model of HOTAIR in a breast cancer model system using the probing reagent 2A3 and an MDA-MB-231 HOTAIR overexpression cell line. We also present a corresponding *in vitro* structure of HOTAIR using SHAPE-MaP with 2A3, enabling direct comparison of structural accessibility between *in cellulo* and *in vitro* contexts and the identification of potential protein binding sites. We also identify candidate RNA modification sites by using a parallel mutational profiling method for modification detection. The structures reported here can serve as a guide for further mechanistic studies of HOTAIR structure-function relationships. Finally, we built large-scale alignments of HOTAIR homologs from a dataset of unannotated primate genomes. These alignments revealed both high conservation of HOTAIR within the primate lineage and covarying base pairs within specific structural motifs. Together, this study supports the conservation of functional structural elements within HOTAIR and provides a foundation for investigating how RNA structure contributes to HOTAIR function.

## RESULTS

### Pipeline for structure probing of a long RNA in breast cancer

To model the *in cellulo* secondary structure of full-length HOTAIR, we adapted the SHAPE-MaP pipeline in a breast cancer system (**Fig. 1A, B**). A major challenge in predicting lncRNA structures in cellular contexts is obtaining adequate material for high-quality chemical probing. Unlike ribosomal RNAs, which are present at up to 10^6^ copies per cell, functional lncRNAs are often expressed at 1 to 1,000 copies per cell (Wu et al. 2021). This limits the number of unique cDNA molecules available for sequencing and robust per-nucleotide mutation rate calculations. Additionally, HOTAIR’s size of 2.1 kb requires adequate sequencing reads to be distributed uniformly across the transcript, further reducing per-nucleotide coverage. Together, low copy number and transcript length make *in cellulo* SHAPE-MaP challenging for lncRNAs.

**Figure 1.**
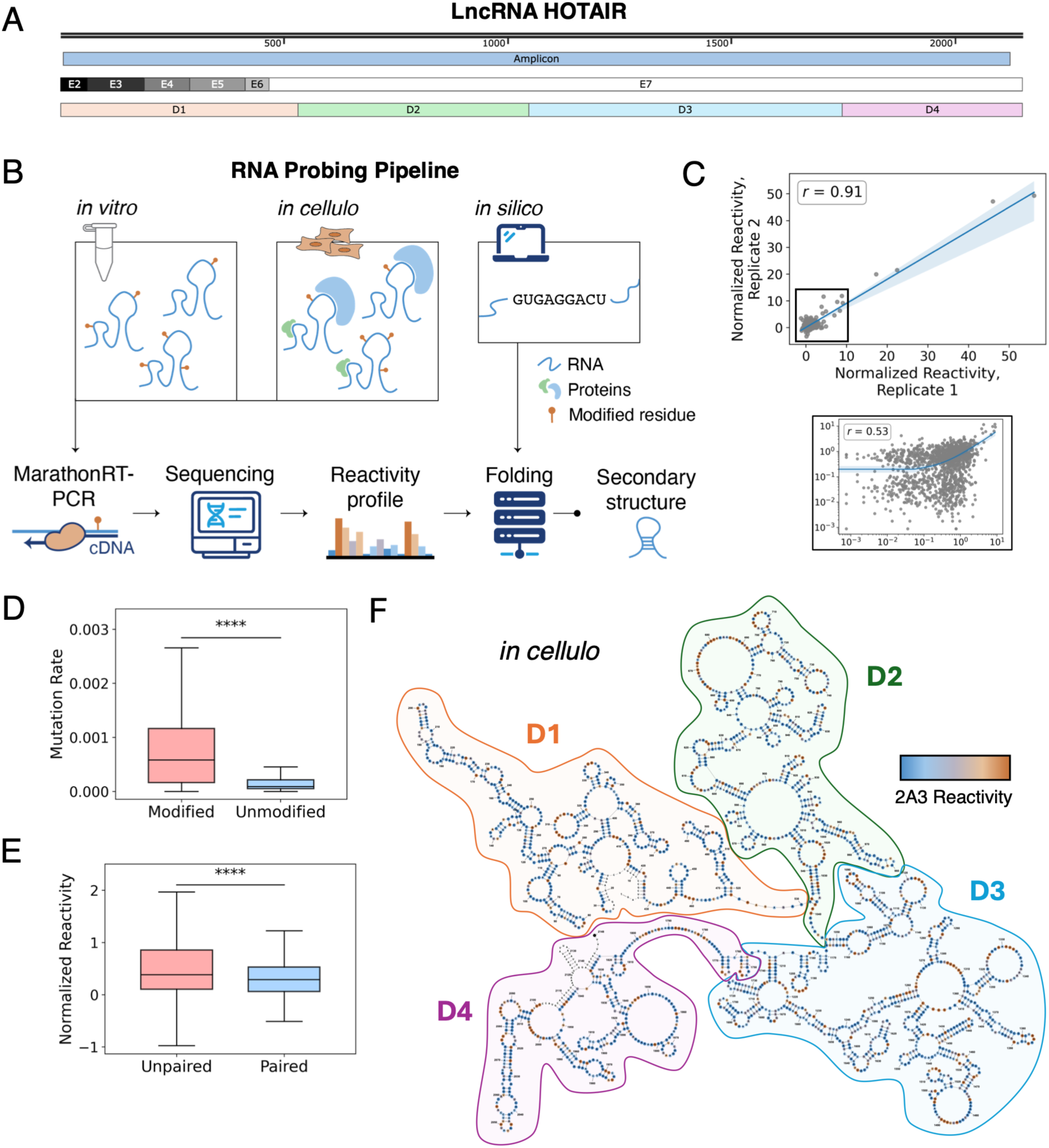
HOTAIR exhibits a complex, multi-domain secondary structure in breast cancer cells. (A) Construct of the lncRNA HOTAIR used in this study. Chemical probing data was captured with a single amplicon spanning 7-2122 nt. Exons are designated from exon 2 to exon 7 (E2-E7), and domains of the *in cellulo* secondary structure are designated as Domain 1 to Domain 4 (D1-D4). Image created using SnapGene. (B) RNA chemical probing pipeline for secondary structure prediction across *in vitro*, *in cellulo*, and *in silico* contexts. Image created using Adobe Stock and Illustrator. (C) Global correlation of normalized reactivities between biological replicates of *in cellulo* SHAPE-MaP on HOTAIR for the full data set or for reactivities < 15. The linear regression line is shown with 95% confidence interval shading. *r*, Pearson’s correlation coefficient. (D) Mutation rates of 2A3-modified samples compared to DMSO-unmodified controls for replicate 1 of *in cellulo* SHAPE-MaP on HOTAIR. Visualized with a box-and-whisker plot. Statistical significance calculated with Mann-Whitney U test; n = 2137 nt (modified), n = 2125 nt (unmodified); **** = p ≤ 0.0001, *** = p ≤ 0.001, ** = p ≤ 0.01. (E) Normalized reactivities for residues predicted to be unpaired compared to paired in the secondary structure prediction generated from replicate-averaged *in cellulo* SHAPE-MaP reactivities of HOTAIR. Visualized with a box-and-whisker plot. Statistical significance calculated with Mann-Whitney U test; n = 975 nt (unpaired), n = 1096 nt (paired); **** = p ≤ 0.0001, *** = p ≤ 0.001, ** = p ≤ 0.01. (F) Secondary structure prediction of HOTAIR using Superfold with replicate-averaged *in cellulo* SHAPE-MaP data. Domains 1-4 are designated as D1-D4. Nucleotides are colored by their normalized reactivity values, orange (≥ 1.0), light orange (≥ 0.75), gray (≥ 0.5), light blue (≥ 0.25), blue ( < 0.25), and white (no data). Structure was visualized with StructureEditor.

Given HOTAIR’s established role in breast cancer progression, we wanted to perform chemical probing in a relevant breast cancer system. Two breast cancer cell lines commonly used in studies of HOTAIR, MDA-MB-231 and MCF7, expressed 0 to 2 copies per cell, respectively, of endogenous HOTAIR, making them unsuitable for chemical probing (**Supplementary Fig. S1A**). To overcome the low copy number problem, we chose an MDA-MB-231 HOTAIR overexpression line (MDA-MB-231 HOTAIR) (Porman et al. 2022), which we found to express HOTAIR at ∼2,030 copies per cell in early-passage cells (passage number 8) but only ∼13 copies per cell in late-passage cells (passage number 20) (**Supplementary Fig. S1B**). Although this overexpression model may not fully reflect endogenous HOTAIR folding, the expression levels of HOTAIR are comparable to those reported in breast cancer patient tissues (Gupta et al. 2010; Arshi et al. 2020). Growing sufficient cells to yield adequate RNA was still challenging due to their slow growth rate and being careful to limit passaging since HOTAIR levels rapidly decrease with increasing passage. Both *in cellulo* SHAPE-MaP replicates were probed at passage number 10, which led to sufficient amplicon generation and sequencing data quality (**Supplementary Table S1**).

Since this cell line had not been used previously for chemical probing, we validated 2A3 probing conditions using 18S rRNA because it is a highly abundant transcript with a well-understood structure. We probed MDA-MB-231 HOTAIR cells with the SHAPE reagent 2A3, which offers enhanced cell-permeability and accuracy compared to other SHAPE reagents (Marinus et al. 2021), with corresponding DMSO-treated negative control samples. Total cellular RNA was extracted and purified, and a ∼1.3 kb region of 18S rRNA was amplified for sequencing and reactivity profile generation via the ShapeMapper2 pipeline (Busan and Weeks 2018). The probing data met sufficient sequencing quality control metrics (**Supplementary Table S1**), and reactivity profiles were highly correlated between two biological replicates (Pearson correlation coefficient, r = 0.997; **Supplementary Fig. S1C**) and significantly higher mutation rates in 2A3-modified 18S rRNA compared to unmodified controls in both replicates (**Supplementary Fig. S1D**).

After validating chemical probing conditions, the same total cellular RNA was used to amplify a single, ∼2.1 kb amplicon spanning 7-2122 nt of HOTAIR. We took advantage of the highly processive reverse transcriptase MarathonRT to allow contiguous amplification of a single sequencing amplicon for *in cellulo* samples, eliminating amplicon-level bias. The MDA-MB-231 HOTAIR cells overexpress the canonical isoform of HOTAIR, which contains exons 2 through 7 (**Fig. 1A**). The probing data for HOTAIR met sufficient sequencing quality control metrics, and over 98% of nucleotides met the minimum read depth of 5,000 reads for each replicate (**Supplementary Table S1, Supplementary Figure S1E**). Reactivity profiles showed high global correlation between two biological replicates (r = 0.91; **Fig. 1C**). Removing the most reactive nucleotides (normalized reactivity > 15) reduced the correlation to weakly positive (r = 0.53). This is consistent with recent *in cellulo* probing studies of lncRNAs, in which lowly reactive nucleotides show increased noise (Smola et al. 2016; Falese et al. 2025). Mutation rates were significantly higher in 2A3-modified samples compared to unmodified controls in both replicates (**Fig. 1D; Supplementary Fig. S1F**), indicating that the reactivity data was sufficient for proceeding to secondary structure prediction. Additional DMSO-treated control samples from SHAPE-MaP were used with MarathonRT in the MRT-ModSeq pipeline to predict candidate modification sites (Tavares et al. 2023). The MDA-MB-231 HOTAIR cell line and technical benefits of MarathonRT thus enabled determination of an *in cellulo* structure and features of HOTAIR in breast cancer.

### Characterizing the global architecture of HOTAIR in cells

Reactivity values were averaged from both biological replicates and used to constrain secondary structure predictions using Superfold (Reuter and Mathews 2010). As expected, nucleotides predicted to be unpaired showed significantly higher reactivity values compared to nucleotides predicted to be paired (**Fig. 1E**), indicating that the average structure is consistent with experimental reactivity profiles in addition to the individual replicates (**Supplementary Fig. S1G**). The secondary structure model revealed that HOTAIR folds into four complex structural domains in breast cancer cells (**Fig. 1F**). Domains were defined as regions comprising complete structures separated by stretches of at least 3 single-stranded nucleotides, and their definition was guided by the prior *in vitro* structure of HOTAIR (**Supplementary Table S2)**. Domain 1 includes exons 2 through 6, while exon 7 spans the end of Domain 1 and extends through Domains 2-4 (**Fig. 1A**).

Notably, the prior *in vitro* study reported that HOTAIR folds into four independent structural domains (Somarowthu et al. 2015). The nucleotide boundaries of Domains 1 and 2 closely match those reported *in vitro*, whereas Domain 3 is longer and more complex *in cellulo*, with a shorter Domain 4. The conservation of overall domain architecture despite the absence of proteins and other cellular factors *in vitro* suggests that HOTAIR folds into a stable structure that is largely determined by its primary sequence. The boundaries of the *in silico* prediction also closely match those *in cellulo*. In contrast, local structural differences, particularly within Domains 3 and 4, may reflect cellular influences on HOTAIR function.

### Identification of highly structured modules in HOTAIR

To enable quantitative comparison of *in cellulo* and *in vitro* models, we predicted a new *in vitro* secondary structure of HOTAIR using SHAPE-MaP with 2A3. Full-length HOTAIR was *in vitro* transcribed, semi-natively purified by size-exclusion chromatography, and refolded at 25 mM Mg^2+^, as described previously (Somarowthu et al. 2015). The probing data passed quality control metrics, including high correlation between biological replicates (r = 0.99) and significantly higher mutation rates in the modified compared to unmodified samples (**Supplementary Fig. S2A, B**). Comparison of replicate-averaged reactivity profiles from *in cellulo* and *in vitro* experiments show overall agreement, with localized regions of increased or suppressed reactivities that may reflect context-dependent differences in RNA accessibility to the probe (**Fig. 2A**).

**Figure 2.**
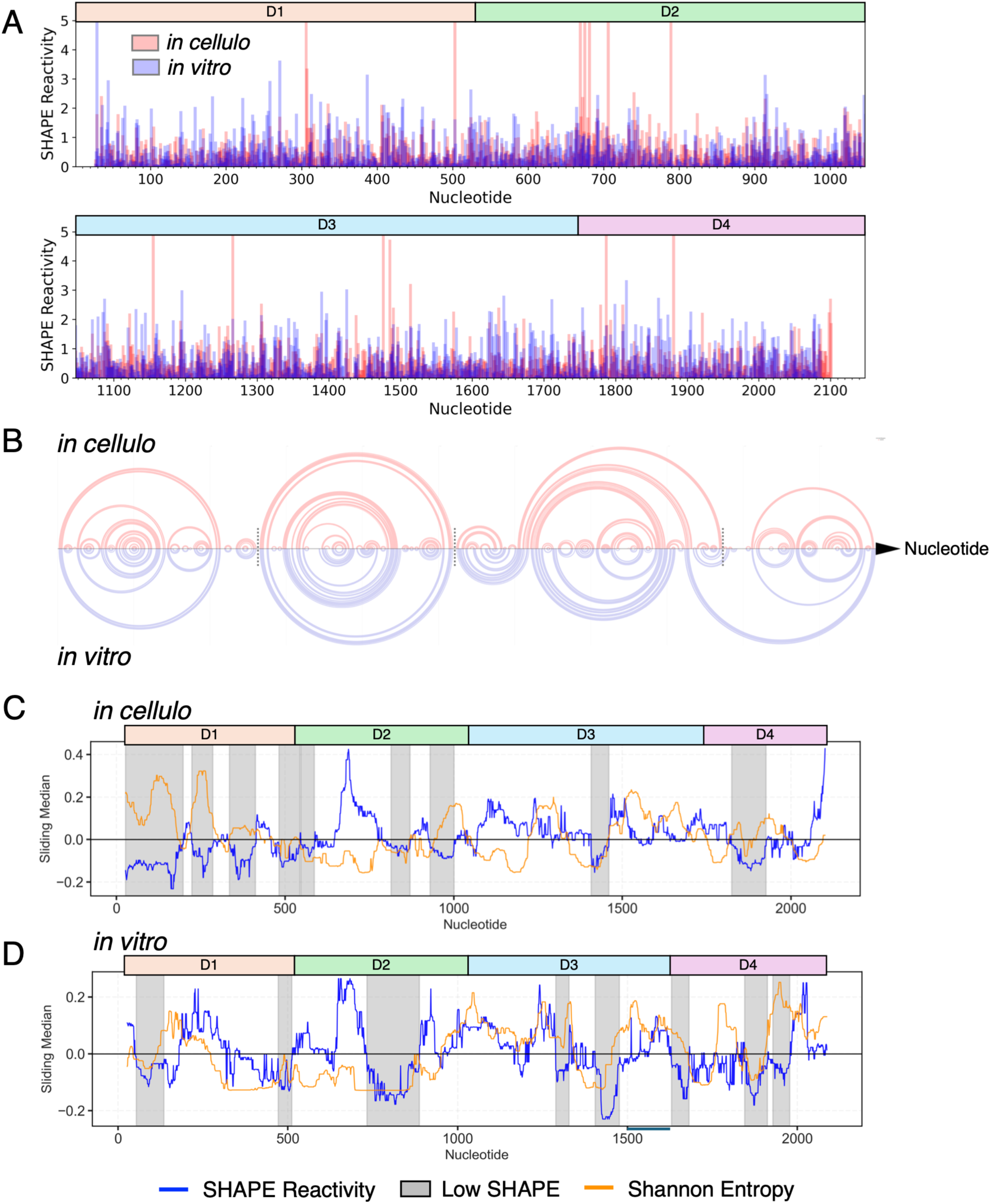
HOTAIR contains highly structured modules across *in cellulo* and *in vitro* contexts. (A) Normalized per-nucleotide reactivities from replicate-averaged *in cellulo* (light red) and *in vitro* (light blue) SHAPE-MaP data, excluding primer binding sites. The range was limited to < 5 to enable comparison of the majority of reactivities. Domains 1-4 of the *in cellulo* structure are designated as D1-D4. (B) Arc plot comparison of the replicate-averaged *in cellulo* and *in vitro* secondary structure predictions. Each arc represents a base pair. Arc plots were created by Rchie. Gray lines represent *in cellulo* domain boundaries. (C) Sliding window median analysis of the replicate-averaged reactivity (blue line) and Shannon entropy (orange line) values relative to the global median for *in cellulo* SHAPE-MaP. Gray boxes indicate regions where the SHAPE reactivity is below a specified threshold for at least 40 nt. Domains 1-4 of the *in cellulo* structure are designated as D1-D4. (D) Sliding window median analysis of the replicate-averaged reactivity and Shannon entropy values for *in vitro* SHAPE-MaP. Visualized as in (C). Domains 1-4 of the *vitro* structure are designated as D1-D4.

Reactivities from individual replicates and replicate-averaged reactivities were used to constrain secondary structure predictions, with higher reactivities mapping to nucleotides predicted to be unpaired in all cases (**Supplementary Fig. S2C, D**). The domain boundaries of the *in vitro* structure inferred in this experiment correspond closely to the previous *in vitro* structure. In this experiment, Domain 3 is predicted to be 115 nt longer, which is closer to that observed *in cellulo* (**Supplementary Table S2**). Despite areas of differing accessibility (**Fig. 2A**), there is substantial shared base-pairing between the *in vitro* and *in cellulo* models, particularly in much of Domains 1 and 2, and the overall four-domain architecture is preserved in both contexts (**Fig. 2B**).

Reactivity and Shannon entropy criteria are commonly used to evaluate the structural stability of motifs within long RNAs. Sliding window median analysis was applied to calculate the replicate-averaged reactivity and entropy profiles relative to the global median (**Fig. 2C, D**). We identified regions of at least 40 continuous nt that have SHAPE reactivity below the global median (< 0), revealing highly structured modules throughout all four domains. Of these highly structured regions, we can categorize them into those with low Shannon entropy (S < 0), suggesting rigid structures, and those with high Shannon entropy (S > 0), suggesting conformationally heterogeneous structures.

Comparison of the base pairing between *in cellulo* and *in vitro* models reveals domain-level differences in HOTAIR folding. Domain 1 is highly structured in both *in cellulo* and *in vitro* contexts, with highly similar base pairing throughout (Jaccard similarity ∼0.8-0.9) (**Supplementary Fig. S2E**). Part of Domain 1 (∼1-350 nt) differs from the *in silico* model showing that our experimental models capture structural information in cells that is different from what is expected from pure thermodynamics. Interestingly, Domain 1 exhibits higher Shannon entropy across much of the domain *in cellulo* (27-182 nt, 224-301 nt, 336-414 nt) compared to *in vitro*.

Domain 2 contains highly structured modules (547-588 nt, 596-645 nt, 788-869 nt) alternating with high reactivity or high Shannon (942-1007 nt) regions *in cellulo*. Both structures have a region of elevated reactivity at ∼700 nt, prior to a low SHAPE, low Shannon region.

In contrast, Domains 3 and 4 contain smaller, highly structured and rigid elements *in cellulo* (1408-1457 nt, 2006-2058 nt). Two low SHAPE regions overlap between the *in cellulo* (1408-1457 nt) and *in vitro* (1405-1453 nt) contexts. Although the overall domain architecture is similar in both contexts, HOTAIR contains several structured modules unique to the cellular context, which may be important to the cellular roles of HOTAIR.

### Candidate molecular interaction sites in HOTAIR

We next identified nucleotide-level differences in reactivities between replicate-averaged *in cellulo* and *in vitro* reactivities using the deltaSHAPE pipeline (Smola et al. 2015a) (**Fig. 3A; Supplementary Fig. S3**). Positive ΔSHAPE values (> 0) indicate nucleotides that are more protected from modification in cells, which may reflect protein or RNA interactions, molecular crowding, or structural stabilization. Negative ΔSHAPE values (< 0) indicate nucleotides that are more accessible in cells, which may reflect sites of structural flexibility or unfolding.

**Figure 3.**
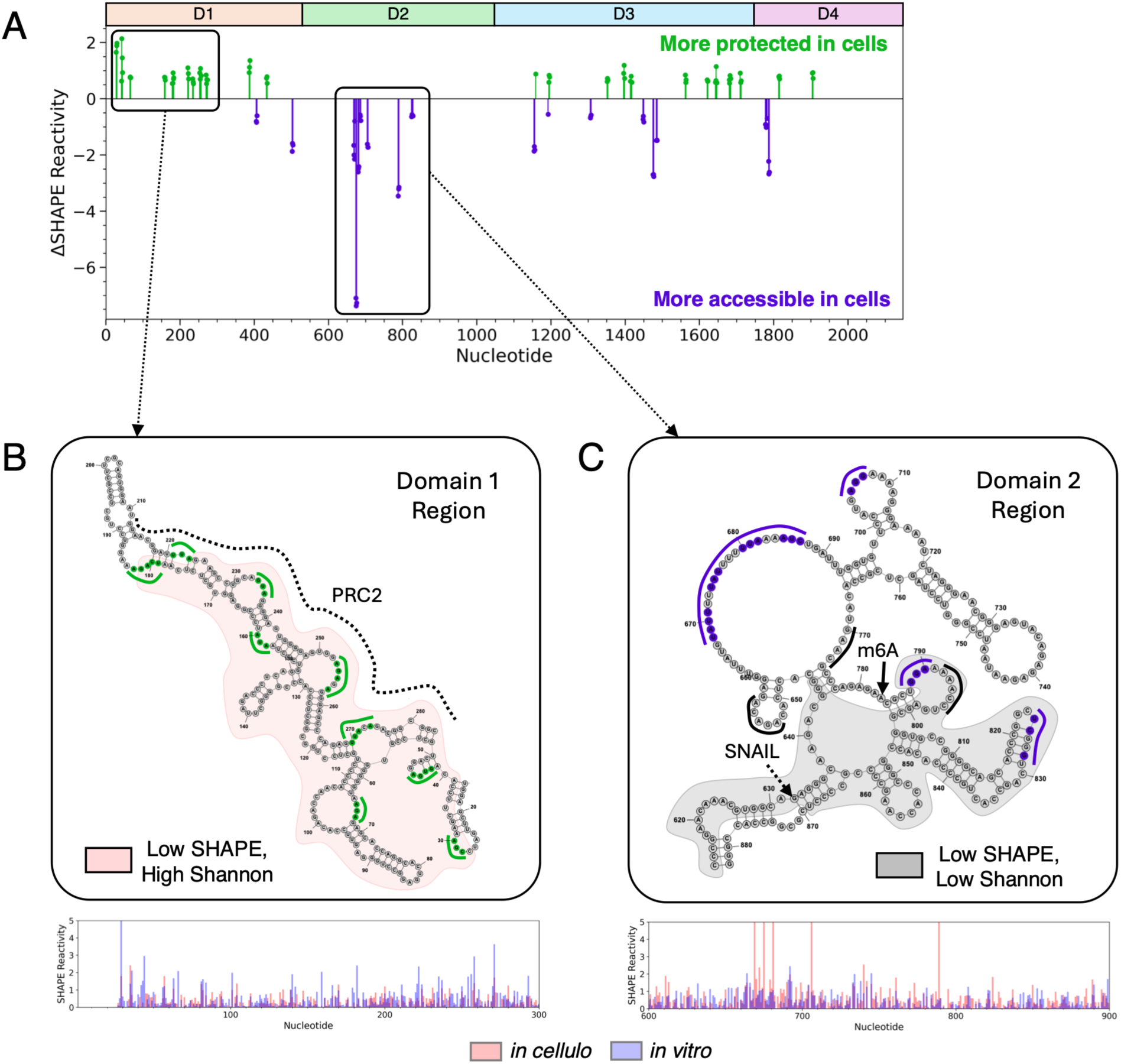
Reactivity differences between *in cellulo* and *in vitro* HOTAIR correlate with sites of molecular interactions. (A) Nucleotides with significant reactivity differences calculated from subtracting replicate-averaged *in cellulo* from *in vitro* reactivities using the deltaSHAPE pipeline. Green indicates sites that are more protected *in cellulo* (ΔSHAPE > 0) and purple indicates sites that are more accessible *in cellulo* (ΔSHAPE < 0). (B) Region of Domain 1 (16-283 nt) with sites of increased protection *in cellulo* (green). This overlaps with a low SHAPE, high Shannon region (light red; 27-182 nt, 224-301 nt) and the minimal PRC2 binding site (black curve; 212-300 nt). Corresponding replicate-averaged reactivity values are shown below the structure. Structure was visualized with StructureEditor. (C) Region of Domain 2 (614-882 nt) with sites of increased accessibility *in cellulo* (purple). This overlaps with a low SHAPE, low Shannon region (gray; 596-645 nt, 788-869 nt), the SNAIL binding site (starting nt indicated with dashed arrow; 632-916 nt), and several m6A sites (black arrow or curve). Corresponding replicate-averaged reactivity values are shown below the structure. Structure was visualized with StructureEditor.

Domain 1 contains eleven sites of increased protection and two sites of increased accessibility in cells, each spanning 3-4 nt. The beginning of Domain 1 (16-283 nt) contains a series of highly structured stems that harbor 9 protected sites. These sites overlap with the previously determined PRC2 minimal binding domain (1-300 nt (Tsai et al. 2010) or 212-300 nt (Wu et al. 2013)) (**Fig. 3B**). Many of these sites are predicted to be single-stranded, suggesting they could remain structurally accessible for intermolecular interactions. This region also displays relatively high Shannon entropy (**Fig. 2C**), although it is unclear if protein binding dynamics are the source of this predicted conformational heterogeneity.

A portion of Domain 2 contains a cluster of seven accessible sites (614-882 nt; **Fig. 3C**) and is flanked by a highly structured, low Shannon entropy region (**Fig. 2C**). This locally stable structure surrounded by a more accessible loop region may permit molecular interactions or tertiary compaction. This region overlaps with a significant portion of the SNAIL binding domain (632-916 nt) and several experimentally mapped N6-methyladenosine (m6A) modification sites (Porman et al. 2022; Garbo et al. 2025). Finally, Domains 3 and 4 have alternating sites of increased protection or accessibility in cells, and seven protected sites overlap with the LSD1 binding site (1500-2146 nt) (Tsai et al. 2010). Overall, many of the deltaSHAPE sites correlate with known HOTAIR protein binders (PRC2 (Tsai et al. 2010; Wu et al. 2013), SNAIL (Garbo et al. 2025), and LSD1 (Tsai et al. 2010)), though we cannot rule out that alternative intermolecular or intramolecular interactions could produce similar protection profiles.

Cellular RNA from unmodified SHAPE-MaP samples can be further subjected to the MRT-ModSeq pipeline to discover candidate sites of natural modification (Tavares et al. 2023). We identified candidate N1-methyladenosine (m1A1243) and N7-methylguanosine (m7G1759) sites in Domain 3 and 4, respectively, which are robustly detected by MRT-ModSeq based on mutation rate filtering **(Supplementary Fig. S3B)**. Sequencing revealed that both sites had significantly elevated mutation rates across all three biological replicates, generally 2-4 orders of magnitude higher than sites predicted to be unmodified. Average Mn²⁺ and Mg²⁺ mutation rates (± SD) were 0.71 ± 0.03 and 0.69 ± 0.05 for m1A1243, and 0.76 ± 0.05 and 0.74 ± 0.05 for m7G1759, respectively. Sequencing confirmed that these sites were not merely genomic DNA mutations **(Supplementary Fig. S3C)**, although orthogonal validation is required to confirm their presence.

### Conservation and covariation of HOTAIR in primates

Although HOTAIR is present across mammals, its sequence rapidly diverges over evolutionary time (He et al. 2011), so we restricted our analysis to the primate lineage. To better assess conservation but still have sufficient diversity for covariation, we analyzed 190 newly sequenced genomes representing 15 of 16 primate families (Kuderna et al. 2023). Since these genomes are unannotated, we identified candidate loci by searching each genome for homologous sequences to human HOTAIR and then extracting the best-scoring hit for each species (**Supplementary Fig. S4A**). We found that searching each genome with the full-length transcript of HOTAIR resulted in poor alignment quality, and in many genomes, only recovered exon 7. Instead, we performed three separate searches with different exon groups (exons 2-4, exons 4-6, and exon 7) against each genome. We performed two validation checks to ensure accurate detection of the HOTAIR locus. First, we confirmed that the best-scoring hits for all exons were on the same scaffold as one another. Next, we validated that these same hits were found on the same scaffold as two genes that neighbor HOTAIR (HOXC11 and HOXC12). With this approach, we validated the HOTAIR locus in 173 out of 190 nonhuman primate genomes. Using the validated sequences, we built a multiple sequence alignment for each exon group, followed by removal of large gaps (> 1,000 nt) not present in human and concatenation of the full locus (**Supplementary Fig. S4A**).

Per-nucleotide conservation was assessed across the full alignment by measuring information content and the frequency of non-gapped positions in sliding windows across HOTAIR (**Fig. 4A**). Regions with high information content and non-gapped frequency reflect positions with high nucleotide conservation and alignment coverage, respectively. From this, we observed that HOTAIR is well-conserved across the analyzed primates, with a median conservation of 1.87/2 bits and median coverage of 98.28%. Though there are several peaks of higher conservation in Domains 1 and 2, all 4 domains are equally well-conserved. We observed some sequence regions in primates that were absent in the human sequence, which may be true indels, misidentified intronic regions, or artifacts of poor sequencing quality.

**Figure 4.**
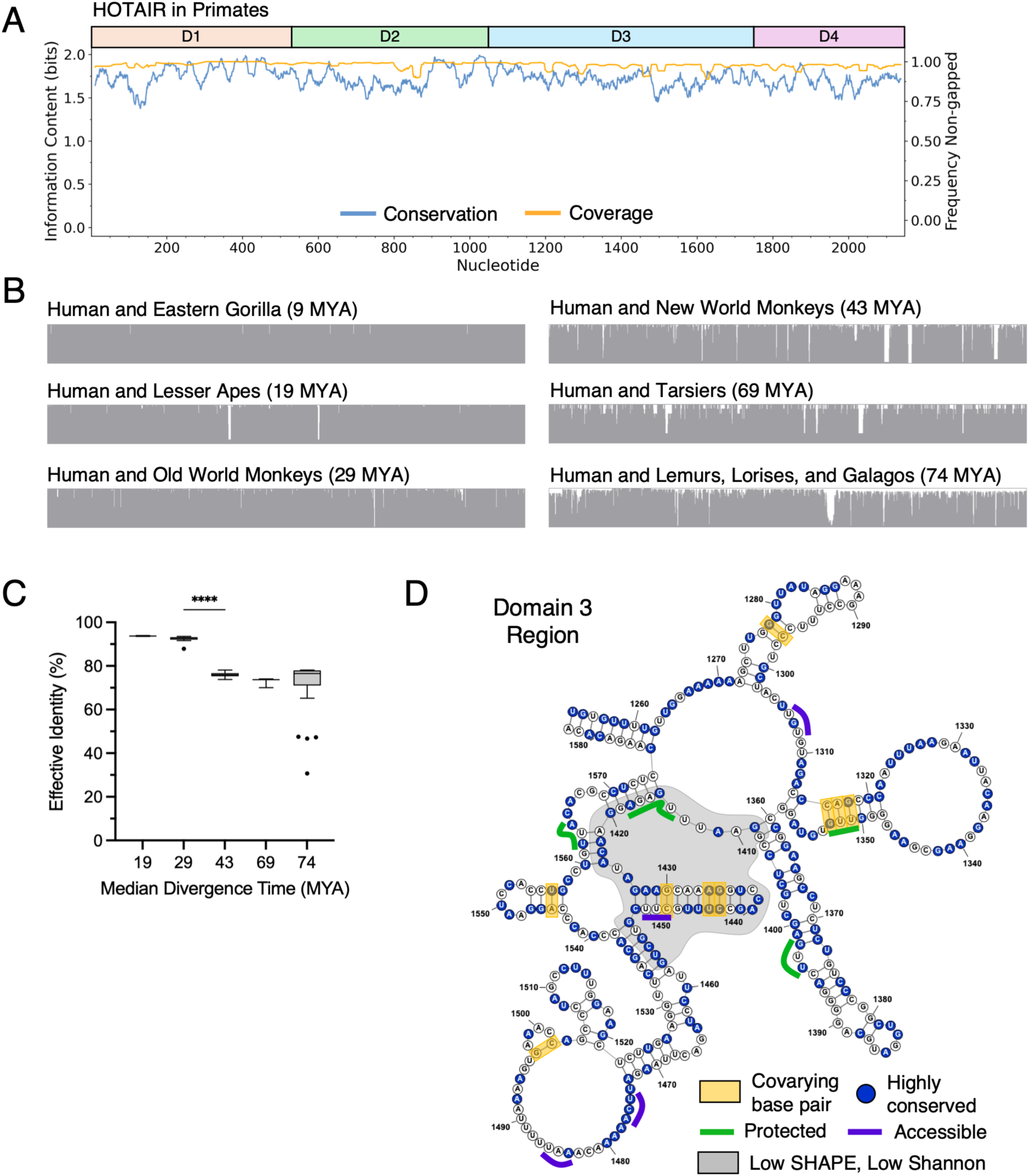
HOTAIR is well-conserved across the primate lineage with evidence for a conserved secondary structure in Domain 3. (A) Rolling average of information content and non-gapped frequency across HOTAIR from a multiple sequence alignment of primates. Domains 1-4 for the *in cellulo* structure are designated as D1-D4. (B) Conservation histograms of subalignments of each primate taxon extracted from the multiple sequence alignment. Gray bars represent conservation for each column in the alignment. Visualized in UGENE. (C) The pairwise effective identity for each primate species compared to human HOTAIR grouped by median divergence time from human in million years ago (MYA). Data are shown as box-and-whisker plots with outliers as individual points. Statistical significance calculated with Kruskal-Wallis test with Benjamini Hochberg correction; n = 8 (19 MYA), n = 54 (29 MYA), n = 70 (43 MYA), n = 3 (69 MYA), n = 37 (74 MYA); **** = p ≤ 0.0001. Plotted in GraphPad Prism. (D) Region of Domain 3 with covarying base pairs (gold), protected sites (green), and accessible sites (purple). Nucleotides with scores > 1.8 bits (the product of information content and non-gapped frequency) from the alignment in (A) are designated as highly conserved. This overlaps with a low SHAPE, low Shannon region (gray; 1408-1457 nt). Structure was visualized with StructureEditor.

To assess HOTAIR conservation over evolutionary time, the primate species were grouped based on established primate clades. A subalignment of each clade and human was extracted from the full alignment and gaps present in all species were removed (**Fig. 4B**). The median percent identity and coverage compared to human for each group was calculated (**Supplementary Table S3**). High coverage across all primate clades indicates good alignment quality. As expected, the percent identities generally decrease with increasing divergence time from human. The effective identity, which scales the identity by the coverage, was calculated between each species and human. This revealed a significant decrease between primate clades that diverged 29 MYA and 43 MYA, which represents Old and New World monkeys, respectively (**Fig. 4C**). Even so, the median effective identity of the most ancient divergence group from 74 MYA, the lemurs, lorises, and galagos, has a median effective identity of 76.51%. However, in the clade just outside the primate lineage (including rodents, rabbits, and tree shrews), HOTAIR is much less conserved across the transcript (**Supplementary Fig. S4B**), with only 50.86% median identity, even though the coverage is > 90% for all species **(Supplementary Table S3**). Although HOTAIR’s sequence is well-maintained across primates, it diverges rapidly after > 85 MYA from human.

This alignment was subsequently filtered to remove similar sequences using a 95% sequence identity threshold, which resulted in 27 representative sequences that were realigned (**Supplementary Fig. S4D**). This alignment was used to build a maximum likelihood tree to predict the phylogenetic relationships purely based on the HOTAIR sequence (**Supplementary Fig. S4E**). The predicted tree successfully recapitulates all primate clades present in this alignment. Additionally, the tree can resolve the lemurs from the lorises and galagos, which are distinct infraorders. We also observe accurate branching of different families within the New World monkeys, showing that HOTAIR can capture more subtle divergence patterns. This further validates our identification of the HOTAIR loci and shows that its sequence does not diverge randomly but tracks with known primate phylogenies.

We next assessed covariation on the 95% clustered alignment, since we hypothesized that the removal of redundantly conserved sequences would capture sufficient sequence diversity to detect compensatory mutations. All covariation was assessed with the RAFSp statistic in sliding windows with E-value < 0.05, an approach which has been optimized for lncRNAs (Tavares et al. 2019). The initial assessment on this alignment predicted 1 covarying base pair in Domain 1, 5 pairs in Domain 3, and 1 pair in Domain 4 (**Supplementary Table S4**). Further covariation analysis on alignments clustered by 98 and 99% sequence identity to include more sequences predicted many of the same base pairs, 2 additional pairs in Domain 3, and 1 additional pair in Domain 4. Regardless of the clustering level, the majority of covarying base pairs were predicted in Domain 3. Thus, we extracted a subalignment of Domain 3 from the full-length, 95% clustered alignment. This identified a maximum of 9 covarying base pairs supported by the *in cellulo* structure, including 3 additional base pairs not predicted with the full-length alignments. In total, we predicted 10 putatively covarying base pairs in Domain 3 (**Supplementary Table S4**). We then visualized the mutational pattern of each of these pair across all 174 primate sequences (**Supplementary Figure S5**). The observed compensatory mutations often included one-sided mutations to GU wobble pairs, but we also detected rarer, two-sided mutations in multiple species (**Supplementary Figure S5i-ix**). This revealed that 1 of the 10 base pairs was merely detected due to high conservation, which is an expected limitation of RAFSp (**Supplementary Figure S5x**) (Tavares et al. 2019). We thus provide support for 9 covarying base pairs consistent with the *in cellulo* secondary structure in Domain 3, suggesting the presence of conserved structures in this domain (**Fig. 4D**). Interestingly, this region overlaps with several protected and accessible sites in cells (**Fig. 3A**) and a low SHAPE, low Shannon region that was identified in both the *in cellulo* and *in vitro* contexts (**Fig. 2C**). An 11-bp stem (1427-1451nt) within the low SHAPE, low Shannon region contains 3 covarying base pairs (**Fig. 4D**). These observations suggest that Domain 3 may contain conserved, stably structured elements with functional relevance.

Overall, these results show that HOTAIR maintains a global four-domain architecture *in cellulo* with local structural and accessibility differences relative to *in vitro* that correspond to known protein binding sites. In addition, we provide evidence for sequence and structural conservation across the primate lineage in HOTAIR.

## DISCUSSION

In this work, we present a full-length SHAPE-constrained *in cellulo* secondary structure model of HOTAIR in the triple negative breast cancer cell line MDA-MB-231 and generated a corresponding *in vitro* structure for quantitative comparison. The structural models predict that HOTAIR adopts a four-domain architecture in both contexts but exhibits domain-specific structural differences in cells. In addition, bioinformatic analyses of newly available primate genomes reveal conservation of the HOTAIR sequence and structural conservation of a specific domain. Together, these results provide a framework for more precise structure-function mechanistic studies of HOTAIR in cellular and disease contexts.

Comparison of the *in cellulo* and *in vitro* models revealed both shared and context-dependent structural features that may influence HOTAIR function in cells. The consistency of the four-domain architecture indicates that the global fold is largely encoded by the primary sequence and can form independently of cellular components, such as protein binding partners, chaperones, or molecular crowding. In contrast, structural differences are observed at the individual domain level, suggesting that cellular factors modulate specific structural regions in HOTAIR. For example, there are several low SHAPE reactivity regions that are unique to the *in cellulo* structure. The boundaries of Domains 3 and 4 also differ slightly between contexts, which may be due to the involvement of protein binders, consistent with the fact that these domains overlap with the reported LSD1 binding site (1500-2148 nt) (Tsai et al. 2010). These observations suggest a general folding principle for long RNAs, where the overall architecture is stably encoded by the sequence, but cellular interactions tune local structural features and domain boundaries. Furthermore, single-stranded ‘linker’ regions that separate the discrete structures contained within each domain may provide flexibility for tertiary folding.

Well-structured modules are typically defined as regions with both low SHAPE reactivity and low Shannon entropy across a defined nucleotide window (Siegfried et al. 2014). However, this approach only captures the most static structures within an RNA. By conducting a parallel analysis that identifies regions of low SHAPE reactivity corresponding to high Shannon entropy, we can identify highly structured regions that may exhibit conformational heterogeneity, consistent with recent approaches (Luo et al. 2025). For example, well-paired RNA structures or switches that sample two or more conformational states could be identified through this approach. Structures of this type are predicted in Domain 1 and 2 *in cellulo*, and regions such as this, which are classified as highly structured but are not necessarily rigid, may represent an important class of RNA motifs for which multiple conformations are essential for function.

Beyond structure determination, directly comparing experimental probing data between *in cellulo* and *in vitro* contexts reveals areas of differential reactivity. Generally, these sites are not evenly interspersed on HOTAIR but tend to form ‘hotspots’ of either increased protection or accessibility. Notably, there is a region in Domain 1 with multiple protected sites that correspond to the reported PRC2 binding domain (1-300 nt (Tsai et al. 2010) or 212-300 nt (Wu et al. 2013)). The structure of a 212-300 nt *in vitro* transcribed fragment was previously predicted using enzymatic mapping, but this prediction differs considerably from the same region in our full-length prediction, highlighting the importance of surrounding sequence context for accurate folding predictions (Wu et al. 2013). However, two protected sites from our study (234-236 nt and 270-274 nt) overlap with previously reported sites protected from nuclease cleavage in the presence of the PRC2 subunits EZH2-EED, suggesting these functional interactions may not depend on the exact base pairing (Wu et al. 2013). Additionally, five protected sites from our study (28-30 nt, 234-236 nt, 253-257 nt, 270-274 nt, and 433-435 nt) overlap with previously reported sites of protection upon JAM2 RNA binding (Balas et al. 2021). This Domain 1 region also overlaps with a low SHAPE, high Shannon region observed *in cellulo*, and we cannot rule out the possibility that other binding interactions may be confounding these measurements. Low SHAPE reactivity may partially reflect protection due to protein or RNA binding rather than a stable structure, while high Shannon entropy may reflect computational uncertainty in the prediction rather than true conformational heterogeneity.

Conversely, the SNAIL binding domain overlaps with sites of increased accessibility in Domain 2, suggesting that these types of sites may also facilitate protein binding. The low SHAPE, low Shannon region here may help fold the structure to create a modular platform for protein binding. Interestingly, the mapped m6A sites in this region all correspond to single-stranded regions, making them accessible to modification-interacting enzymes. This region also exhibits distinct base pairing from the *in silico* and *in vitro* models (**Supplementary Fig. S2E**), making it an interesting structural target unique to the cellular context. We also provide evidence for novel modifications in Domains 3 and 4, which opens new possibilities for HOTAIR-protein regulation. Overall, HOTAIR’s structure is consistent with protein interactions that promote epigenetic silencing and cancer progression. We hope that these findings provide a structural foundation to guide future studies of the HOTAIR-protein interactome in cancer.

Because lncRNAs often diverge rapidly in sequence, restricting conservation analyses to primates allows for better assessment of conservation. Homologous HOTAIR sequences were identified across a diverse dataset of 190 primate genomes, although incomplete assemblies and unannotated genomes hampered identification of the HOTAIR loci in all cases. Scaffold validation with neighboring genes on the *HOXC* locus confirmed correct locus identification in 173 unannotated primates, which still provided sufficient genomic data to reveal substantial sequence conservation across the entire HOTAIR transcript.

Within the primate lineage, we analyzed HOTAIR divergence across evolutionary time from 19 MYA to 74 MYA, with a significant divergence event between 29 MYA and 43 MYA. The 19 MYA and 29 MYA divergence groups represent Hylobatidae (lesser apes) and Cercopithecidae (Old World monkeys), respectively, while the 43 MYA, 69 MYA, and 74 MYA groups represent Platyrrhini (New World monkeys), Tarsiidae (tarsiers), and Strepsirrhini (lemurs, lorises, and galagos). Changes in conservation across these clades may partly reflect the influence of geographic dispersal on HOTAIR sequence divergence.

Furthermore, a representative alignment of HOTAIR can accurately recapitulate known primate phylogenies with family-level resolution, reminiscent of a molecular clock. While multiple rRNA genes have historically been required to define phylogenetic relationships, a single lncRNA achieves comparable resolution here. This suggests that HOTAIR, and potentially other lncRNAs, may be reliable indicators of evolutionary divergence, at least within primates.

Taking advantage of the primate genomes allowed inclusion of novel species and provided sufficient phylogenetic diversity for covariation analysis. We show evidence for covarying base pairs in Domain 3, supporting the functional relevance of a discrete structural module here. We also capture subtle compensatory base pair mutations across primate evolution. This element overlaps with a low SHAPE reactivity and low Shannon entropy region and is close to several protected and accessible sites, all of which indicate potential functional relevance. Only a few covarying base pairs were predicted in Domains 1 and 4, although our alignments may not have sufficient sequence diversity to detect additional covariation. However, by limiting our analysis to primate genomes, we show that HOTAIR is far more conserved than generally appreciated for a lncRNA and provide evidence that selective pressure may act on specific structural elements.

A limitation of this study is that HOTAIR overexpression was required to obtain sufficient transcript abundance for probing. Although the overexpression levels are comparable to some patient tumors, the structure presented here may not fully recapitulate native folding in the tumor environment. Future studies that probe endogenous HOTAIR, such as in patient-derived xenograft cell lines (Finlay-Schultz et al. 2020) or primary breast tumor samples (Gupta et al. 2010), would help address this limitation. Additionally, chemical probing reports indirectly on RNA structure and captures an ensemble-averaged snapshot, rather than distinct conformational states. RNAs, like proteins, likely adopt multiple conformations, some of which may guide distinct cellular functions. Future studies in which libraries are constructed from specific subcellular compartments can address this issue. Finally, identification and alignment of lncRNAs across species remains challenging due to frequent insertions and deletions, exon variation, and incomplete genome assemblies, which limits our ability to detect covariation. Addressing these challenges will allow future studies to make critical links between structure and function in HOTAIR and other lncRNA targets.

The *in cellulo* secondary structure model of HOTAIR presented here provides a roadmap for future mechanistic studies. Regions that are highly structured and show increased protection, conservation, or covariation represent promising candidates for investigating protein binding and the role of HOTAIR’s structure in gene regulation and cancer. More broadly, determining structures of lncRNAs in relevant cellular contexts is a key step for connecting structure to function.

## MATERIALS AND METHODS

### Tissue culture

MCF7 (ATCC HTB-22) and MDA-MB-231 (ATCC HTB-26) cells were obtained from the American Type Culture Collection. MDA-MB-231 pBABE-puro HOTAIR overexpression cells (MDA-MB-231 HOTAIR) had been previously generated from parental MDA-MB-231 cells using retroviral transduction as described (Meredith et al. 2016; Porman et al. 2022). Cells were grown in Dulbecco’s Modified Eagle Medium with high glucose and L-glutamine without sodium pyruvate (GenClone, Gibco) with 10% fetal bovine serum (Avantor) and 1% penicillin-streptomycin (Gibco) at 37°C and 5% CO_2_. Media for MDA-MB-231 HOTAIR cells was supplemented with 1 µg/mL puromycin (Gibco).

### *In vitro* transcription and purification for qPCR

#### In vitro transcription

HOTAIR was transcribed from a pCARs08 plasmid containing the HOTAIR sequence immediately downstream of a T7 promoter. The plasmid had been previously generated as described (Somarowthu et al. 2015). Plasmid was linearized by incubating with SalI restriction enzyme and CutSmart Buffer at 37°C for 16 hrs, followed by enzyme inactivation at 95°C for 5-10 min. Plasmid was purified with the QIAquick Gel Extraction Kit (Qiagen) and added to a 1 mL transcription reaction with transcription buffer (15 mM MgCl2, 40 mM Tris-Cl pH 8, 2 mM spermidine, 10 mM NaCl, 0.01% Triton 100), 10 mM DTT, 4 mM of each NTP at pH 7, RNase OUT (Invitrogen), and 0.08 mg/mL T7 enzyme (purified *in house*). Reaction was incubated at 37°C for 2 hrs, followed by plasmid digestion with 4 µL TURBO DNase (Invitrogen) at 37°C for 30 min and addition of 10 µl of 0.5 M Na_2_EDTA pH 8.5. Transcribed RNA was concentrated to 500 µL and exchanged into 150 mM KCl and 50 mM MOPS pH7 using a 100 kDa Amicon Ultra 0.5 mL filtration column (Amicon).

#### Gel purification

Urea loading dye (11 M urea, 2% xylene cyanol/bromophenol blue, 50% sucrose, 40 mM Tris-HCl pH 7.5, 0.8 mM Na_2_EDTA pH 8.5 pH 8) was heated at 65°C for 10 min and added to the buffer-exchanged RNA in a 1:1 ratio. An 8 M Urea-5% Acrylamide-Tris-Borate-EDTA (TBE) denaturing gel was cast onto custom-made siliconized gel plates from the Yale University Scientific Glassblowing Laboratory and polymerized with ammonium persulfate and TEMED. The gel was run at 20 W for 30 min prior to loading the RNA sample, which was run at 20-25 W for 3 hours. Gel plates were opened and plastic wrap was laid on top of the gel to flip onto a gel tray. RNA was then visualized under UV light, and the band was extracted with a sterile blade. The RNA was eluted with the ‘crush and soak’ method. Briefly, the gel slice was forced through a 5 ml syringe, 10-12 mL of elution buffer (300 mM NaCl, 10 mM MOPS pH 6, 1 mM Na_2_EDTA pH 8.5) was added, and RNA was eluted at 4°C overnight with shaking. Gel pieces were removed with a filter. 3 M sodium-acetate pH 5.2 was added to precipitate the RNA at –80°C for 2 hrs. To obtain a pellet, the sample was spun at 15,000g for 30 min, washed with 70% EtOH, and spun at 15,000g for another 30 min. The pellet was allowed to dry and redissolved in 1x ME storage buffer (10 mM MOPS pH 6.5, 1 mM Na_2_EDTA pH 8.5). Concentration was determined by Nanodrop at A260.

### Reverse transcription quantitative polymerase chain reaction (RT-qPCR)

#### Reverse transcription (RT)

To generate a standard curve of *in vitro* transcribed HOTAIR, a 10-fold serial dilution of 10^11^ copies/µL to 10^3^ copies/µL was made using the molecular weight calculated from AAT Bioquest (AAT Bioquest 2026). For total cellular RNA samples, cells were dislodged with cell scrapers, centrifuged, and resuspended in Dulbecco’s PBS (GenClone, Gibco). RNA was extracted using 3 parts TRIzol (Invitrogen) to 1 part cell suspension followed by adding chloroform:isoamyl alcohol 24:1 (Sigma) equal to 1/5 TRIzol volume. After centrifuging at 3,000g for 15 min at 4°C, the aqueous RNA phase was removed and precipitated in 75% final EtOH at –80°C for several hours. RNA was pelleted at 15,000g for 30 min at 4°C, air dried, and resuspended in 1x ME buffer. Samples were treated with DNase I (Qiagen) prior to clean-up with the RNeasy kit (Qiagen). For annealing, 1 µg of each RNA sample was combined with 10 mM dNTP Mix (NEB), 2 mM RT primer, and RNase-free water and incubated at 65°C for 5 min in a PCR thermocycler, then placed on ice for 1 min. To the annealing mix, 5x First Strand Synthesis buffer (Invitrogen), 10 mM DTT, 1 µL SUPERase•In RNase inhibitor (Invitrogen), and 1 µL Superscript (SS) II or III (Invitrogen) were added for a final RT reaction volume of 20 µL. Reverse transcription proceeded at 55°C for 50 min for SSIII or 42°C for 50 min for SSII followed by enzyme inactivation at 85°C for 5 min. Template RNA was degraded via alkaline hydrolysis in which 1 µL 3 M KOH was added, the solution was incubated at 95°C for 5 min and 4°C for 2 min, and then 1 µL 3 M HCl was added. The cDNA was cleaned with 1.8x AMPure XP beads (Beckman) according to the manufacturer’s protocol.

#### Quantitative polymerase chain reaction (qPCR)

For each qPCR reaction, 5 µL cleaned cDNA, 2x Light Cycler 480 SYBR Green I Master Mix (Roche), 10 µM forward primer, 10 µM reverse primer, and RNase-free water was combined for a 20 µL reaction. The qPCR plate was covered with a plastic sheet and centrifuged at 2,000g for 2 min. Plates were immediately placed in the qPCR thermocycler and subjected to pre-incubation, amplification (40 cycles), melting curve, and cooling steps.

#### qPCR analysis

Technical duplicates and/or biological replicates were averaged. The standard curve was plotted and fit to a linear regression line. The slope and y-intercept were used to determine the copies/µl of HOTAIR in cellular samples and converted to copies/cell based on the volume of RNA per dish and the cell count determined using a hemocytometer and Trypan Blue (Sigma). All primers for qPCR are listed in **Table of Primers**.

### *In cellulo* SHAPE probing

#### Harvesting cells

MDA-MB-231 HOTAIR cells were grown to 80-90% confluency on two 150 mm dishes per biological replicate. The media was aspirated, the cells were washed with phosphate-buffered saline (PBS), and 10 mL PBS was added to each dish. Cells were dislodged with a cell scraper, pooled, and centrifuged at 1,000g for 5 min at 4°C. Cell pellets were resuspended in 1.5 mL PBS, then evenly split into treated and control fractions.

#### In cellulo SHAPE probing

Solution of 1 M 2A3 was prepared immediately prior to probing by dissolving 100 mg of 2A3 into 530 µL DMSO (Sigma) and protected from light. For the 2A3-modified fraction, 100 µL of 1 M 2A3 was added per 0.5 mL cell suspension for a final concentration of 100 mM and mixed well. For the DMSO-unmodified control fraction, 100 μL of 100% DMSO was added per 0.5 mL cell suspension and mixed well. Both fractions were incubated at 37°C for 15 minutes protected from light.

#### Total cellular RNA extraction

To quench the reaction, 3 parts TRIzol (Invitrogen) were added to 1 part cell suspension, mixed well, and incubated for 5 min at room temperature. Next, chloroform:isoamyl alcohol 24:1 (Sigma) equal to 1/5 TRIzol volume was added, mixed well, and incubated for 5 min at room temperature. Samples were centrifuged at 4,700 rpm for 15 min at 4°C, and the aqueous RNA phase was precipitated with 1 µL glycogen per 1 mL aqueous RNA and 75% final EtOH at –20°C for 2-3 days.

#### EtOH precipitation and RNA cleanup

RNA was pelleted by spinning at 15,000g for 30 min at 4°C. After air drying, each RNA pellet was resuspended in RNase-free water, and in-solution DNase digestion was performed with TURBO DNase buffer and 5 µL TURBO DNase (Invitrogen) at 37°C for 30 min. DNase-treated RNA was purified using the RNeasy kit (Qiagen) and eluted in 1x ME buffer (10 mM MOPS pH 6.5, 1 mM Na_2_EDTA pH 8.5) supplemented with SUPERase•In RNase inhibitor (Invitrogen). Concentration was determined by Nanodrop at A260.

### *In vitro* SHAPE probing

#### In vitro transcription

HOTAIR pCARs08 plasmid was linearized with SalI-HF restriction enzyme (NEB) and CutSmart Buffer at 37°C for 1 hr with no heat inactivation. Plasmid was precipitated with 1/10 volume of 3 M NaOAc pH 5.5 and 75% EtOH at –20°C for 1 hr. Plasmid was pelleted by spinning at 21,000g for 30 min at 4°C, washed with 70% EtOH, and spun at 21,000g for 5 min. Pellet was allowed to air dry before resuspending in RNase-free water. *In vitro* transcription was performed with transcription buffer (15 mM MgCl2, 40 mM Tris-Cl pH 8, 2 mM spermidine, 10 mM NaCl, 0.01% Triton 100), 10 mM DTT, 2.5 mM of each NTP at pH 7, SUPERase•In RNase inhibitor (Invitrogen), thermostable inorganic pyrophosphatase (NEB), 10 mM MgCl_2_ (Invitrogen) and 0.2 mg/mL T7 enzyme (purified *in house*) for a 1 mL reaction. Reaction was incubated at 30°C for 1.5 hrs, followed by plasmid digestion with 20 µL TURBO DNase (Invitrogen) at 37°C for 10 min and addition of 20 µl of 0.5 M Na_2_EDTA pH 8.5. Transcribed RNA was concentrated and exchanged into filtration buffer (50 mM K-HEPES pH 7.2, 150 mM KCl, 0.1 mM Na_2_EDTA pH 8.5) with 100 kDa Amicon Ultra 0.5 ml filtration columns (Amicon).

#### Semi-native purification and folding

RNA was semi-natively purified with a fast performance liquid chromatography AKTA using a Sephacryl S-400 column (Cytiva) equilibrated with filtration buffer. RNA elution into 0.5 mL fractions was monitored using the absorbance at A260. The peak fraction was collected, and concentration was determined with Nanodrop at A260. For folding, MgCl_2_ was added to a final concentration of 25 mM and incubated at 37°C for 30 min.

#### In vitro SHAPE probing

Immediately after folding, the sample was split into 5 µg fractions for each probing condition and replicate. To modified fractions, 2A3 was added for a final concentration of 100 mM. An equivalent volume of DMSO (Sigma) was added to unmodified control fractions. Both fractions were mixed well and incubated at 37°C for 10 min protected from light. To quench the reaction, 1/10 volume of 3 M NaOAc and 75% EtOH were added to each sample and precipitated at –20°C overnight. To pellet RNA, samples were spun at 21,100g for 30 min at 4°C, washed with 70% EtOH, and spun at 21,100g for 5 min at 4°C. Residual EtOH was removed, and the pellet was air dried and resuspended in RNase-free water. Concentration was determined by Nanodrop at A260.

### RT-PCR, library preparation, and sequencing

#### RT-PCR

All RNA samples from *in cellulo* and *in vitro* SHAPE probing were reverse transcribed with MarathonRT (Guo et al. 2020). Annealing mixes were prepared with 1 µg RNA and 1 µM HOTAIR-specific (or 18S rRNA-specific) primer and annealed at 68°C for 5 min. To each reaction, 2.5x MarathonRT buffer (125 mM Tris-HCl pH 7.5, 500 mM KCl, 12.5 mM DTT, 1.25 mM dNTPs (NEB, Thermo), 2.5 mM MnCl_2_ (Sigma)), MarathonRT (purified *in house*), and glycerol were added and incubated at 42°C for 3 hours, followed by enzyme inactivation at 70°C for 15 min. Single-stranded cDNA was cleaned with either 1.8x RNAClean XP or 1.8x AMPure XP beads (Beckman). Each amplicon was amplified with PCR using Q5 High-Fidelity 2X MasterMix (NEB), 10 µM forward primer, 10 µM reverse primer, and 5 µL cleaned cDNA with touchdown PCR conditions. Annealing temperatures were estimated using SnapGene or the NEB Tm Calculator. PCR products were cleaned with 1.8X AMPure XP beads (Beckman). For *in cellulo* samples, HOTAIR was prepared as a single amplicon (7-2122 nt), while for *in vitro* samples, HOTAIR was tiled with two overlapping amplicons (7-1213 nt and 1010-2122 nt). The control 18S rRNA was prepared as a single amplicon (32-1304 nt) from the same total cellular RNA used for HOTAIR. All primers for SHAPE-MaP are listed in **Table of Primers**.

#### Library preparation and sequencing

DNA concentration was measured with the Qubit dsDNA HS Assay Kit (Invitrogen) and diluted to 0.2 ng/µL. Tagmentation and Nextera PCR were carried out according to the Nextera XT DNA Library Prep documentation (Illumina), except at 1/5 the volume. Libraries were purified with 1.8X AMPure XP beads (Beckman) and analyzed on a Bioanalyzer with the High Sensitivity DNA Kit or TapeStation with the HS D5000 ScreenTape Assay (Agilent). Final libraries were adjusted to 4 nM, pooled based on the desired read depth, and adjusted to the final loading concentration required for each kit. Sequencing was performed with the NextSeq 1000/2000 P1 Reagents Kit or the NextSeq 1000/2000 P1 XLEAP-SBS Reagent Kit (Illumina) on a NextSeq 1000/2000.

### Sequencing alignment and structure prediction

Sequencing files were retrieved using the BaseSpace Sequence Hub Downloader and transferred to an HPC cluster. Samples with two amplicons were merged prior to alignment. Paired-end reads were aligned to the full-length HOTAIR sequence, excluding nucleotides up to and including the primer binding sites, using ShapeMapper2 under default parameters (Busan and Weeks 2018). After verifying that samples met sufficient quality control thresholds, the average reactivity per nucleotide was calculated between two biological replicates to create.map files for secondary structure prediction with SuperFold under default parameters (Reuter and Mathews 2010). The *in silico* structure prediction was carried out identically, except the SHAPE reactivity values in the.map file were set to –999 to indicate no experimental data.

### Identification of structured regions

The rolling median of normalized SHAPE reactivity (from ShapeMapper2) and Shannon entropy (from SuperFold) values was calculated in sliding windows of 51 nt around the center nucleotide with step size of 1 nt and truncated edge windows. The global median was subtracted from each rolled value. Typically, low SHAPE regions are defined by reactivity less than the global median (< 0), but we set an arbitrary threshold of < –0.02 to exclude noisy regions near the global median. We identified all low SHAPE regions < –0.02 for a minimum length of 40 nt. We further classified these into low SHAPE and low Shannon entropy (< 0), indicating highly structured regions that adopt a dominant conformation, and low SHAPE and high Shannon entropy (> 0), since these regions are highly structured yet may be conformationally heterogeneous.

### Jaccard similarity calculation

The Jaccard similarity was calculated between each pair of structures in sliding windows of 301 nt around the center nucleotide with step size of 1 nt and truncated edge windows. The Jaccard similarity is defined as the number of shared base pairs (intersection) divided by the total number of unique base pairs (union) in each window.

### ΔSHAPE analysis

Replicate-averaged *in cellulo* SHAPE reactivities were subtracted from replicate-averaged *in vitro* SHAPE reactivities using the deltaSHAPE pipeline under default parameters and statistical thresholds (Smola et al. 2015a). The original deltaSHAPE.py script was first converted to Python 3 compatibility.

### MRT-ModSeq pipeline

The same DMSO-treated control samples from *in cellulo* SHAPE-MaP were further subjected to the MRT-ModSeq pipeline (Tavares et al. 2023). Briefly, reverse transcription was performed with MarathonRT in the presence of 1 mM Mg^2+^ or 2 mM Mn^2+^. Samples were amplified with two overlapping amplicons (7-1114 nt and 1010-2122 nt) and sequenced according to **RT-PCR, library preparation, and sequencing**. The two amplicons were merged and aligned to HOTAIR with ShapeMapper2, with Mg^2+^ libraries designated as “untreated” and Mn^2+^ libraries designated as “modified.” Mutation rate filtering was used to identify the presence of candidate RNA modifications using the established criteria, where m1A sites have a characteristic Mn^2+^ mutation rate > 0.5 and m7G sites have a characteristic Mn^2+^ mutation rate > 0.05 and Mg^2+^ mutation rate > 0.0015 (Tavares et al. 2023). Only modifications passing these criteria and present in three biological replicates are reported.

Genomic DNA (gDNA) from MDA-MB-231 HOTAIR cells was extracted with the Monarch Spin gDNA Extraction Kit. A specific region of the gDNA containing potential modifications was amplified with PCR (see **Table of Primers**) and samples were sent for Sanger sequencing. Other modifications were identified via machine learning, including an average of 34 candidate pseudouridine (Ψ) sites per replicate, but these sites tend to have high false positive rates, and only 8 Ψ sites were present in at least two biological replicates. Furthermore, the sequences surrounding the candidate Ψ sites did not match any known pseudouridine synthase recognition motifs, so these modifications were discarded.

### HOTAIR annotation pipeline

#### Exon-aware searches of unannotated genomes

Assembled primate genomes were downloaded from the European Nucleotide Archive (Accession PRJEB67744), previously deposited by Kuderna et al (Kuderna et al. 2023). For each genome, the tool Exonerate (version 2.4.0) with the est2genome model (Slater and Birney 2005) was used to find potential matches to the given query sequence. Since searches with full-length human HOTAIR (hHOTAIR) returned sequences with poor coverage, three independent searches were conducted with exons 2-4 (288 nt), exons 4-6 (279 nt), and exon 7 (1683 nt). Multiple exons were combined where necessary to reduce false positives from searching with short exons. For each search, the hit with the best alignment score was extracted and converted to RNA using a custom script. Due to the computational requirements, searches were conducted in parallel on a high-performance computing cluster.

#### Scaffold validation

Correct identification of HOTAIR sequences was validated by checking that all exons were present on the same scaffold. This served as initial validation that our searches produced hits from the same genomic locus. Secondly, we validated that the neighboring genes HOXC11 and HOXC12 were on the same scaffold as the exons of HOTAIR. All of exon 1 of HOXC11 (NM_014212.4) and the last 500 bases of exon 2 of HOXC12 (NM_173860.3) were identified in each genome with Exonerate and used as references. Of the 190 unannotated primate species, HOTAIR sequences from 173 nonhuman primate species were successfully validated.

#### Multiple sequence alignment and locus construction

The final set of sequences and hHOTAIR (174 sequences total) were aligned using MAFFT (version 7.526, –-auto flag) (Katoh and Standley 2013). Inspection of these initial multiple sequence alignments revealed that human exons 2-6 successfully aligned across primates but were separated by large intervening regions (>1,000 nt) absent in humans. These may represent introns that were misidentified by Exonerate, and these regions were removed from each alignment. Finally, the three separate alignments were concatenated to generate the full locus.

### Building a mammalian multiple sequence alignment

To assess HOTAIR conservation outside the primate lineage, HOTAIR mammalian alignments were obtained from the MultiZ 100-way track in the UCSC Genome Browser, then filtered with Galaxy (Galaxy 2026) to only retain species within the Scandentia order and Glires clade. Sequences from the 100-way alignment were extracted, the human HOTAIR sequence used in this study was inserted, and the HOTAIR sequences from selected species were re-aligned with MAFFT (Katoh and Standley 2013). Due to the inclusion of introns in the 100-way alignment, columns in the alignment where there was a gap present in the human HOTAIR sequence were removed with the Biostrings library in R (https://bioconductor.org/packages/Biostrings).

### Conservation and evolutionary divergence analysis

The Esl-alistat package in HMMER (version 3.4, http://hmmer.org/) was used to calculate per-column information content in bits and non-gapped frequency in each input alignment. Per-nucleotide scores for hHOTAIR were extracted, and the rolling average was plotted in sliding windows of 21 nt around the center nucleotide with step size of 1 nt. For pairwise conservation analysis, the percent identity and coverage of each species compared to human was calculated using the NCBI MSA Viewer (version 1.26.0) after setting hHOTAIR as the anchor sequence. We define ‘effective identity’ as the product of percent identity and coverage to account for sequences with lower coverage.

For evolutionary divergence analysis, each primate was binned by its median pairwise divergence time from human, which was retrieved from TimeTree (Kumar et al. 2022). For divergence bins with at least 3 species, the distribution of effective identities was analyzed. Subalignments of each primate taxon and hHOTAIR were extracted from the 180-species multiple sequence alignment, and gaps present in all species were removed. To compare outside the primate lineage, hHOTAIR was aligned to annotated mouse HOTAIR (NCBI Reference Sequence NR_047528.1) using Clustal Omega (Madeira et al. 2024), and percent identity and coverage were calculated with the NCBI MSA Viewer. All alignments and conservation histograms were visualized with UGENE (version 52.0).

### Clustering and phylogeny analysis

To reduce redundancy from similar sequences, the full alignment (180 sequences) was filtered by 95, 98, or 99% sequence identity (–c flag) using CD-HIT-EST (version 4.8.1) with a word size of 8 (–n flag) (Li and Godzik 2006; Fu et al. 2012). This resulted in 28, 41, and 60 sequences, respectively. After ensuring hHOTAIR was included as the representative sequence of its cluster, they were realigned with MAFFT (version 7.526, –-auto flag). Clustered alignments were used for covariation analysis, as described below. The phylogenetic tree was constructed from the 95% clustered alignment using the PhyML Maximum Likelihood method in UGENE (version 52.0). The General Time Reversible (GTR+Γ) substitution model with 4 substitution rate categories was used to allow for unequal base frequencies, variable substitution rates, and rate variation across sites. Branch support was assessed with 1,000 bootstrap replicates in UGENE. The tree was unrooted and visualized in MEGA with the radial layout (version 12.1.1).

### Covariation analysis

To prepare alignments for covariation analysis, input alignments were converted to Stockholm format using the Esl-reformat package in HMMER (version 3.4, http://hmmer.org/). The replicate-averaged *in cellulo* secondary structure was added to the alignment, with gaps inserted as single-stranded positions corresponding to the gapped hHOTAIR sequence. Covarying base pairs were then predicted using R-scape (version 0.2.1) installed from Bioconda (https://anaconda.org/bioconda/rscape/files/manage?page=2) (Rivas et al. 2017). The average product-corrected (APC) RNAalifold Stacking test (–-RAFSp) was selected for its ability to detect subtle covariation in lncRNA alignments, although we also tried the APC G-test (–-GTp) (Woolf 1957; Lindgreen et al. 2006; Dunn et al. 2008; Tavares et al. 2019). Due to the length of HOTAIR, covariation was assessed in sliding windows of 300 or 500 nt (–-window) with step size of 100 nt (–-slide). All base pairs with E-value < 0.05 that were present in the input structure were reported as covarying. Column positions output by R-scape were converted to the correct nucleotide positions in the hHOTAIR sequence. To visualize the identity of the compensatory mutations, each hHOTAIR nucleotide position was converted to the column position in the full-length, pre-clustered alignment of 174 sequences. The column identity (A, U, G, C, or gap) was plotted across the sequences to analyze the covariation patterns, and any base pair that had no evidence of compensatory mutations, but was detected merely due to high conservation, was removed, which was the case for one of the ten predicted covarying base pairs in Domain 3.

### Plotting and statistical analysis

Unless otherwise noted, all plots were generated using Matplotlib or Seaborn in Jupyter Notebook or JupyterLab. For box-and-whisker plots, boxes indicate the interquartile range (IQR), the center line is the median, and the whiskers extend to the farthest data point within 1.5xIQR. Unless otherwise indicated, statistical significance was calculated with a two-tailed Mann-Whitney U test or an unpaired, two-tailed Student’s t-test, where **** = p ≤ 0.0001, *** = p ≤ 0.001, ** = p ≤ 0.01, and * = p ≤ 0.05.

## DATA DEPOSITION

Raw and processed sequencing data from SHAPE-MaP and MRT-ModSeq have been deposited in NCBI’s Gene Expression Omnibus (GEO Accession: GSE331520) (Edgar et al. 2002). Processed SHAPE-MaP, deltaSHAPE, MRT-ModSeq, and bioinformatics data are available as Supplemental Data files. Structure files, reference sequences, multiple sequence alignment files, and analysis code is available on GitHub at https://github.com/pylelab/HOTAIR.

## Supporting information

Supplemental_Data_1_HOTAIR_qPCR

Supplemental_Data_2_HOTAIR_SHAPE-MaP

Supplemental_Data_3_HOTAIR_deltaSHAPE

Supplemental_Data_4_HOTAIR_MRT-ModSeq

Supplemental_Data_5_HOTAIR_Conservation

Supplemental_Data_6_HOTAIR_Covariation

## ACKNOWLEDGEMENTS

We thank Dr. Allison M. Porman Swain and Dr. Aaron M. Johnson for generously gifting the MDA-MB-231 HOTAIR cells that enabled *in cellulo* probing. We thank Dr. Li-Tao Guo for providing the pCARs08 HOTAIR plasmid for *in vitro* work. We thank Dr. Michael Van Zandt and New England Discovery Partners (NEDP) for providing the 2A3 SHAPE-MaP reagent. We thank all former and current members of the Pyle Lab for thoughtful discussions, including Dr. Ananth Kumar, Dr. Rafael Tavares, Dr. Shivali Patel, Dr. Han Wan, Dr. Tanja Hann, Michelle Luo, Lucille Tsao, and Jack Sung. Thank you to the Yale Center for Research Computing for their guidance and use of the HPC clusters. A.M.P. is an investigator of the Howard Hughes Medical Institute. This work was also supported by the NIH (1R01HG011868-01 to A.M.P.), the NSF GRFP (DGE-2139841 to Z.R.P.), and the Yale Hahn Scholars Fellowship (to L.L.).

