## Supplemental_Material for "Structural features and conservation of the lncRNA HOTAIR in primates"

**Supplementary Materials and Methods……………………………………………………3-6**

Human HOTAIR Sequence

Table of Primers

**Supplementary Figures……………………………….……….………………...7-16**

**Supplementary Figure S1.** Validation of SHAPE-MaP chemical probing of HOTAIR and 18S rRNA in breast cancer cells.

**Supplementary Figure S2.** Prediction of SHAPE-constrained *in vitro* secondary structure model of HOTAIR and structure comparison across contexts.

**Supplementary Figure S3.** Differential reactivity sites and candidate chemical modification sites on *in cellulo* HOTAIR.

**Supplementary Figure S4.** HOTAIR annotation and conservation in mammals and primates.

**Supplementary Figure S5.** Compensatory mutations observed in predicted base pairs in HOTAIR Domain 3 across primates.

**Supplementary Tables…………………………………………………………........17-22**

**Supplementary Table S1.** Sequencing quality control metrics of samples from SHAPE-MaP and MRT-ModSeq.

**Supplementary Table S2.** Domain boundaries for *in cellulo*, *in vitro*, and *in silico* HOTAIR secondary structure models.

**Supplementary Table S3.** Conservation of HOTAIR between human and different taxa.

**Supplementary Table S4.** Covarying base pairs detected in Domain 3 from primate alignments of HOTAIR.

**Supplementary References…………………………………………………………..........23**

### Supplementary Materials and Methods

#### Human HOTAIR Sequence

The human HOTAIR sequence used in this study comprises nucleotides 1-2148 (excluding the polyA tail) from GenBank Accession DQ926657.1, deposited by Rinn et al. (Rinn et al. 2007). Exon boundaries were determined from NCBI Reference Sequence NR_003716.4; note that this reference is missing the first 5 nt and has a slightly longer final exon compared to the original HOTAIR sequence. Exon 2, 1-60 nt; Exon 3, 61-186 nt; Exon 4, 187-288 nt; Exon 5, 289-412 nt; Exon 6, 413-465 nt; Exon 7, 466-2148 nt. Exon 1 is present in some annotated isoforms and is separated by a relatively long intron.

GACUCGCCUGUGCUCUGGAGCUUGAUCCGAAAGCUUCCACAGUGAGGACUGCUC

CGUGGGGGUAAGAGAGCACCAGGCACUGAGGCCUGGGAGUUCCACAGACCAACA

CCCCUGCUCCUGGCGGCUCCCACCCGGGGCUUAGACCCUCAGGUCCCUAAUAUCC

CGGAGGUGCUCUCAAUCAGAAAGGUCCUGCUCCGCUUCGCAGUGGAAUGGAACG

GAUUUAGAAGCCUGCAGUAGGGGAGUGGGGAGUGGAGAGAGGGAGCCCAGAGU

UACAGACGGCGGCGAGAGGAAGGAGGGGCGUCUUUAUUUUUUUAAGGCCCCAAA

GAGUCUGAUGUUUACAAGACCAGAAAUGCCACGGCCGCGUCCUGGCAGAGAAAA

GGCUGAAAUGGAGGACCGGCGCCUUCCUUAUAAGUAUGCACAUUGGCGAGAGAA

UUAAGUGCUGCAACCUAAACCAGCAAUUACACCCAAGCUCGUUGGGGCCUAAGC

CAGUACCGACCUGGUAGAAAAAGCAACCACGAAGCUAGAGAGAGAGCCAGAGGA

GGGAAGAGAGCGCCAGACGAAGGUGAAAGCGAACCACGCAGAGAAAUGCAGGCA

AGGGAGCAAGGCGGCAGUUCCCGGAACAAACGUGGCAGAGGGCAAGACGGGCAC

UCACAGACAGAGGUUUAUGUAUUUUUAUUUUUUAAAAUCUGAUUUGGUGUUCC

AUGAGGAAAAGGGAAAAUCUAGGGAACGGGAGUACAGAGAGAAUAAUCCGGGU
CCUAGCUCGCCACAUGAACGCCCAGAGAACGCUGGAAAAACCUGAGCGGGUGCC
GGGGCAGCACCCGGCUCGGGUCAGCCACUGCCCCACACCGGGCCCACCAAGCCCC
GCCCCUCGCGGCCACCGGGGCUUCCUUGCUCUUCUUAUCAUCUCCAUCUUUAUG
AUGAGGCUUGUUAACAAGACCAGAGAGCUGGCCAAGCACCUCUAUCUCAGCCGC
GCCCGCUCAGCCGAGCAGCGGUCGGUGGGGGACUGGGAGGCGCUAAUUAAUUG
AUUCCUUUGGACUGUAAAAUAUGGCGGCGUCUACACGGAACCCAUGGACUCAUA
AACAAUAUAUCUGUUGGGCGUGAGUGCACUGUCUCUCAAAUAAUUUUUCCAUAG
GCAAAUGUCAGAGGGUUCUGGAUUUUUAGUUGCUAAGGAAAGAUCCAAAUGGG
ACCAAUUUUAGGAGGCCCAAACAGAGUCCGUUCAGUGUCAGAAAAUGCUUCCCC
AAAGGGUUGGCAGUGUGUUUUGUUGGAAAAAAGCUUGGGUUAUAGGAAAGCCU
UUCCCUGCUACUUGUGUAGACCCAGCCCAAUUUAAGAAUUACAAGGAAGCGAAG
GGGUUGUGUAGGCCGGAAGCCUCUCUGUCCCGGCUGGAUGCAGGGGACUUGAGC
UGCUCCGGAAUUUGAGAGGAACAUAGAAGCAAAGGUCCAGCCUUUGCUUCGUGC
UGAUUCCUAGACUUAAGAUUCAAAAACAAAUUUUUAAAAGUGAAACCAGCCCUA
GCCUUUGGAAGCUCUUGAAGGUUCAGCACCCACCCAGGAAUCCACCUGCCUGUU
ACACGCCUCUCCAAGACACAGUGGCACCGCUUUUCUAACUGGCAGCACAGAGCA
ACUCUAUAAUAUGCUUAUAUUAGGUCUAGAAGAAUGCAUCUUGAGACACAUGG
GUAACCUAAUUAUAUAAUGCUUGUUCCAUACAGGAGUGAUUAUGCAGUGGGAC
CCUGCUGCAAACGGGACUUUGCACUCUAAAUAUAGGCCCCAGCUUGGGACAAAA
GUUGCAGUAGAAAAAUAGACAUAGGAGAACACUUAAAUAAGUGAUGCAUGUAG
ACACAGAAGGGGUAUUUAAAAGACAGAAAUAAUAGAAGUACAGAAGAACAGAA
AAAAAAUCAGCAGAUGGAGAUUACCAUUCCCAAUGCCUGAACUUCCUCCUGCUA

UUAAGAUUGCUAGAGAAUUGUGUCUUAAACAGUUCAUGAACCCAGAAGAACGC
AAUUUCAAUGUAUUUAGUACACACACAGUAUGUAUAUAAACACAACUCACAGAA
UAUAUUUUCCAUACAUUGGGUAGGUAUGCACUUUGUGUAUAUAUAAUAAUGUA
UUUUCCAUGCAGUUUUAAAAUGUAGAUAUAUUAAUAUCUGGAUGCAUUUUC

**Table of Primers**

| **Primer** | **Sequence (5’ to 3’)** |
|---|---|
| **HOTAIR, qPCR** | |
| qPCR RT | TTCTACCAGGTCGGTACTGG |
| qPCR F (Porman et al. 2022) | TCTGGAGCTTGATCCGAAAG |
| qPCR R (Porman et al. 2022) | GGTGTTGGTCTGTGGAACT |
| **HOTAIR, *in vitro* SHAPE-MaP** | |
| HOTAIR SHAPE-MaP RT Amp 1 | CGGACTCTGTTTGGGCCTCCT |
| HOTAIR SHAPE-MaP PCR F Amp 1 | CCTGTGCTCTGGAGCTTGAT |
| HOTAIR SHAPE-MaP PCR R Amp 1 | ACTCTGTTTGGGCCTCCT |
| HOTAIR SHAPE-MaP RT Amp 2 | ATCTACATTTTAAAACTGC |
| HOTAIR SHAPE-MaP PCR F Amp 2 | AGGCGCTAATTAATTGATTCC |
| HOTAIR SHAPE-MaP PCR R, Amp 2 | CTACATTTTAAAACTGCATGG |
| **HOTAIR, *in cellulo* SHAPE-MaP** | |
| HOTAIR SHAPE-MaP RT | ATCTACATTTTAAAACTGC |
| HOTAIR SHAPE-MaP PCR F | CCTGTGCTCTGGAGCTTGAT |
| HOTAIR SHAPE-MaP PCR R | CTACATTTTAAAACTGCATGG |
| **HOTAIR, *in cellulo* MRT-ModSeq** | |

| | |
|---|---|
| ModSeq RT Amp 1 | TGAGAGACAGTGCACTCAC |
| ModSeq PCR F Amp 1 | CCTGTGCTCTGGAGCTTGAT |
| ModSeq PCR R Amp 1 | GAGACAGTGCACTCACGC |
| ModSeq RT Amp 2 | ATCTACATTTTAAAACTGC |
| ModSeq PCR F Amp 2 | AGGCGCTAATTAATTGATTCC |
| ModSeq PCR R Amp 2 | CTACATTTTAAAACTGCATGG |
| **18S rRNA, *in cellulo* SHAPE-MaP** | |
| 18S SHAPE-MaP RT | GAAAGAGCTATCAATCTGTC |
| 18S SHAPE-MaP PCR F | TGTCTCAAAGATTAAGCCATGC |
| 18S SHAPE-MaP PCR R | AGCTATCAATCTGTCAATCC |
| **gDNA Sequencing** | |
| PCR Forward gDNA | TGTAAAACGACGGCCAGTGGAAAGATCC AAATGGGACC |
| PCR Reverse gDNA | CAGGAAACAGCTATGACCAATTGCGTTC TTCTGGGTTC |

RT, reverse transcription; F, forward; R, reverse; Amp, amplicon

#### Supplementary Figures

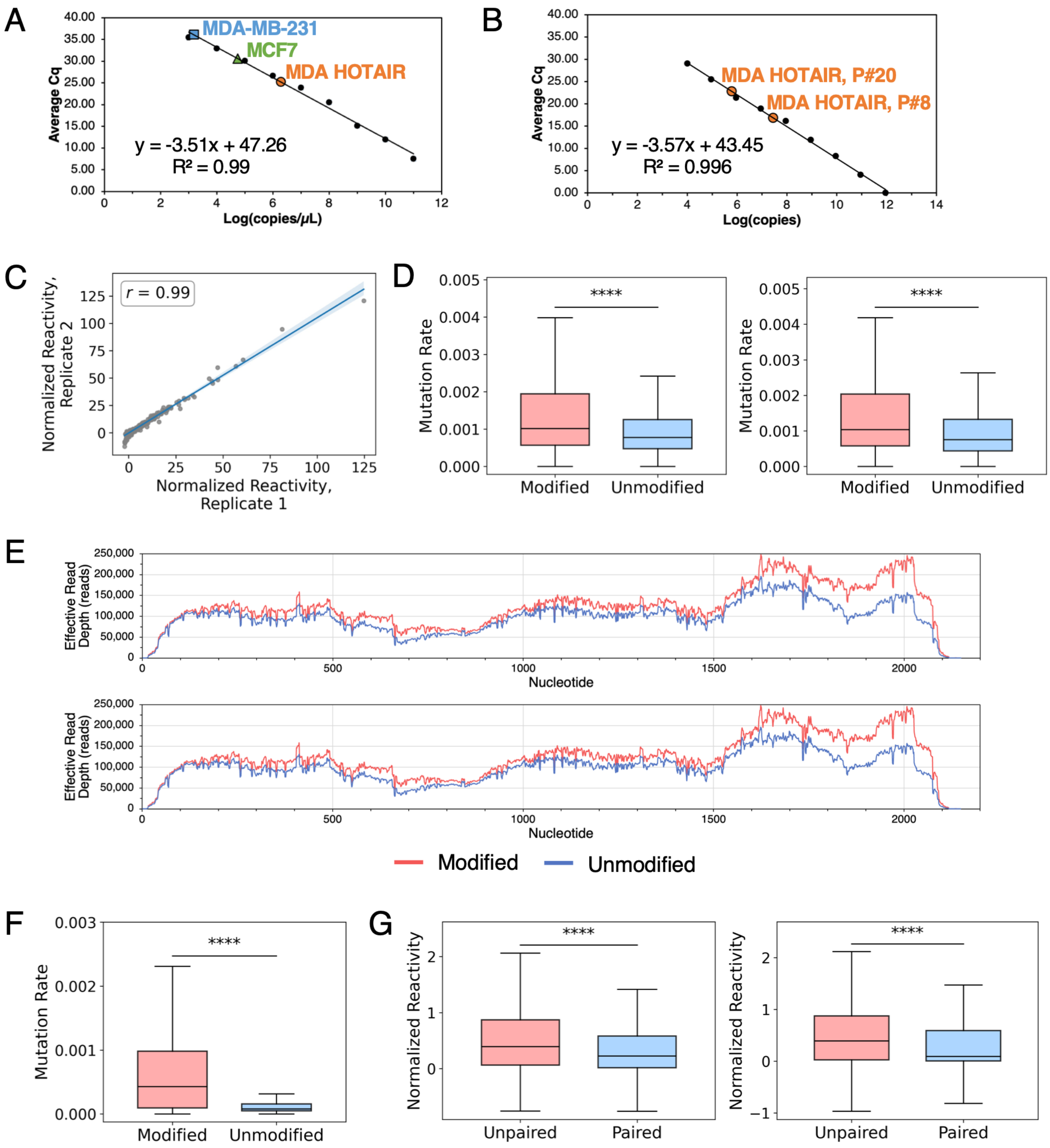


**Supplementary Figure S1.** Validation of SHAPE-MaP chemical probing of HOTAIR and 18S rRNA in breast cancer cells. (A) Expression of HOTAIR in MDA-MB-231, MCF7, and MDA-MB-231 HOTAIR cell lines determined by absolute RT-qPCR. I*n vitro* transcribed HOTAIR was used to generate a standard curve and fitted to a linear regression line to determine copy number. Quantification cycle (Cq) is average of two biological replicates done in duplicate. Plotted in Microsoft Excel. (B) Expression of HOTAIR in MDA-MB-231 HOTAIR lines at two different passage numbers (P#) determined by absolute RT-qPCR. I*n vitro* transcribed HOTAIR was used to generate a standard curve and fitted to a linear regression line to determine copy number. Cq measured in duplicate. Plotted in Microsoft Excel. (C) Global correlation of normalized reactivities between biological replicates of *in cellulo* SHAPE-MaP on18S rRNA. The linear regression line is shown with 95% confidence interval shading. *r*, Pearson's correlation coefficient. (D) Mutation rates of 2A3-modified samples compared to DMSO-unmodified controls for replicate 1 (left) and replicate 2 (right) of *in cellulo* SHAPE-MaP on 18S rRNA. Visualized with a box-and-whisker plot. Statistical significance calculated with Mann-Whitney U test; replicate 1, n = 1279 nt (modified), n = 1304 nt (unmodified); replicate 2, n = 1283 nt (modified), n = 1300 nt (unmodified); **** = $p \leq 0.0001$, *** = $p \leq 0.001$, ** = $p \leq 0.01$. (E) Effective read depth per nucleotide from *in cellulo* SHAPE-MaP on HOTAIR for replicate 1 (top) and replicate 2 (bottom). (F) Mutation rates of 2A3-modified samples compared to DMSO-unmodified controls for replicate 2 of *in cellulo* SHAPE-MaP on HOTAIR. Visualized with a box-and-whisker plot. Statistical significance calculated with Mann-Whitney U test; n = 2134 nt (modified), n = 2134 nt (unmodified); **** = $p \leq 0.0001$, *** = $p \leq 0.001$, ** = $p \leq 0.01$. (G) Normalized reactivities for residues predicted to be unpaired compared to paired in the secondary structure prediction generated from *in cellulo* SHAPE-MaP reactivities of HOTAIR for replicate 1 (left) and replicate 2 (right). Visualized with a box-and-whisker plot. Statistical significance calculated with Mann-Whitney U test; replicate 1, n = 961 nt (unpaired), n = 1110 nt (paired); replicate 2, n = 931 nt (unpaired), n = 1140 nt (paired); **** = $p \leq 0.0001$, *** = $p \leq 0.001$, ** = $p \leq 0.01$.

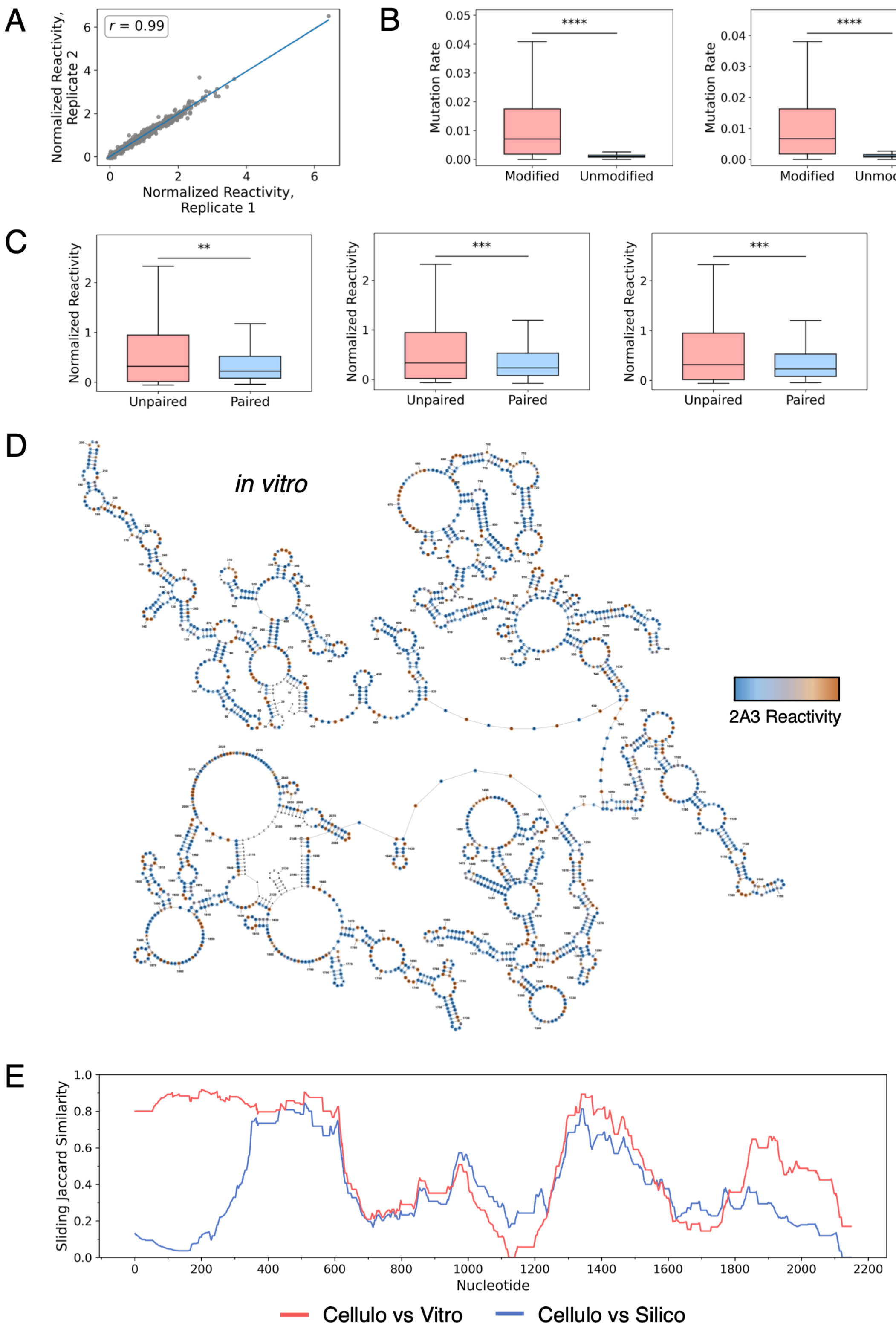


**Supplementary Figure S2.** Prediction of SHAPE-constrained *in vitro* secondary structure model of HOTAIR and structure comparison across contexts. (A) Global correlation of normalized reactivities between biological replicates of *in vitro* SHAPE-MaP on HOTAIR. The linear regression line is shown with 95% confidence interval shading. *r*, Pearson's correlation coefficient. (B) Mutation rates of 2A3-modified samples compared to DMSO-unmodified controls for replicate 1 (left) and replicate 2 (right) of *in vitro* SHAPE-MaP on HOTAIR. Visualized with a box-and-whisker plot. Statistical significance calculated with Mann-Whitney U test; replicate 1, n = 2128 nt (modified), n = 2148 nt (unmodified); replicate 2, n = 2136 nt (modified), n = 2148 nt (unmodified); **** = $p \leq 0.0001$, *** = $p \leq 0.001$, ** = $p \leq 0.01$. (C) Normalized reactivities for residues predicted to be unpaired compared to paired in the secondary structure prediction generated from *in vitro* SHAPE-MaP reactivities of HOTAIR from replicate 1 (left), replicate 2 (middle), and the replicate-averaged values (right). Visualized with a box-and-whisker plot. Statistical significance calculated with Mann-Whitney U test; replicate 1, n = 963 nt (unpaired), n = 1095 nt (paired); replicate 2, n = 963 nt (unpaired), n = 1098 nt (paired); average, n = 954 nt (unpaired), n = 1104 nt (paired); **** = $p \leq 0.0001$, *** = $p \leq 0.001$, ** = $p \leq 0.01$. (D) Secondary structure prediction of HOTAIR using Superfold with replicate-averaged *in vitro* SHAPE-MaP data. Nucleotides are colored by their normalized reactivity values, orange (≥ 1.0), light orange (≥ 0.75), gray (≥ 0.5), light blue (≥ 0.25), blue ( < 0.25), and white (no data). Structure was visualized with StructureEditor. (E) Jaccard similarity comparisons in sliding windows for the *in cellulo*, *in vitro*, and *in silico* secondary structure models of HOTAIR.

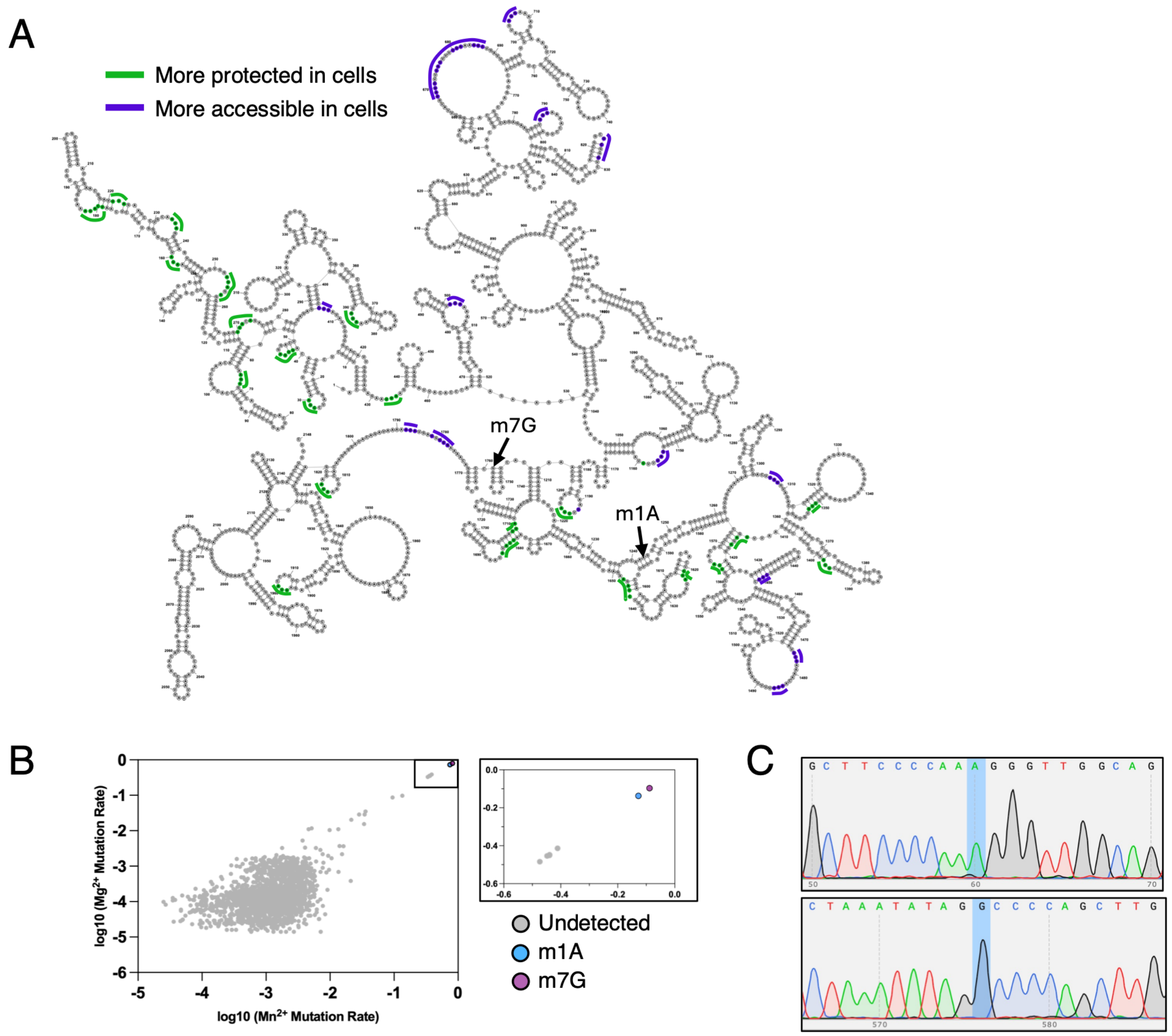


**Supplementary Figure S3.** Differential reactivity sites and candidate chemical modification sites on *in cellulo* HOTAIR. (A) Structure of full-length HOTAIR *in cellulo* with sites that are more protected *in cellulo* (green, ΔSHAPE > 0) and sites that are more accessible *in cellulo* (purple, ΔSHAPE < 0). Structure was visualized with StructureEditor. (B) Representative replicate from MRT-ModSeq. Plotted in GraphPad Prism. The m1A and m7G modification sites identified here are shown on the structure in (A). (C) Sequencing of genomic DNA for region corresponding to the m1A and m7G sites identified in (B). Visualized in SnapGene.

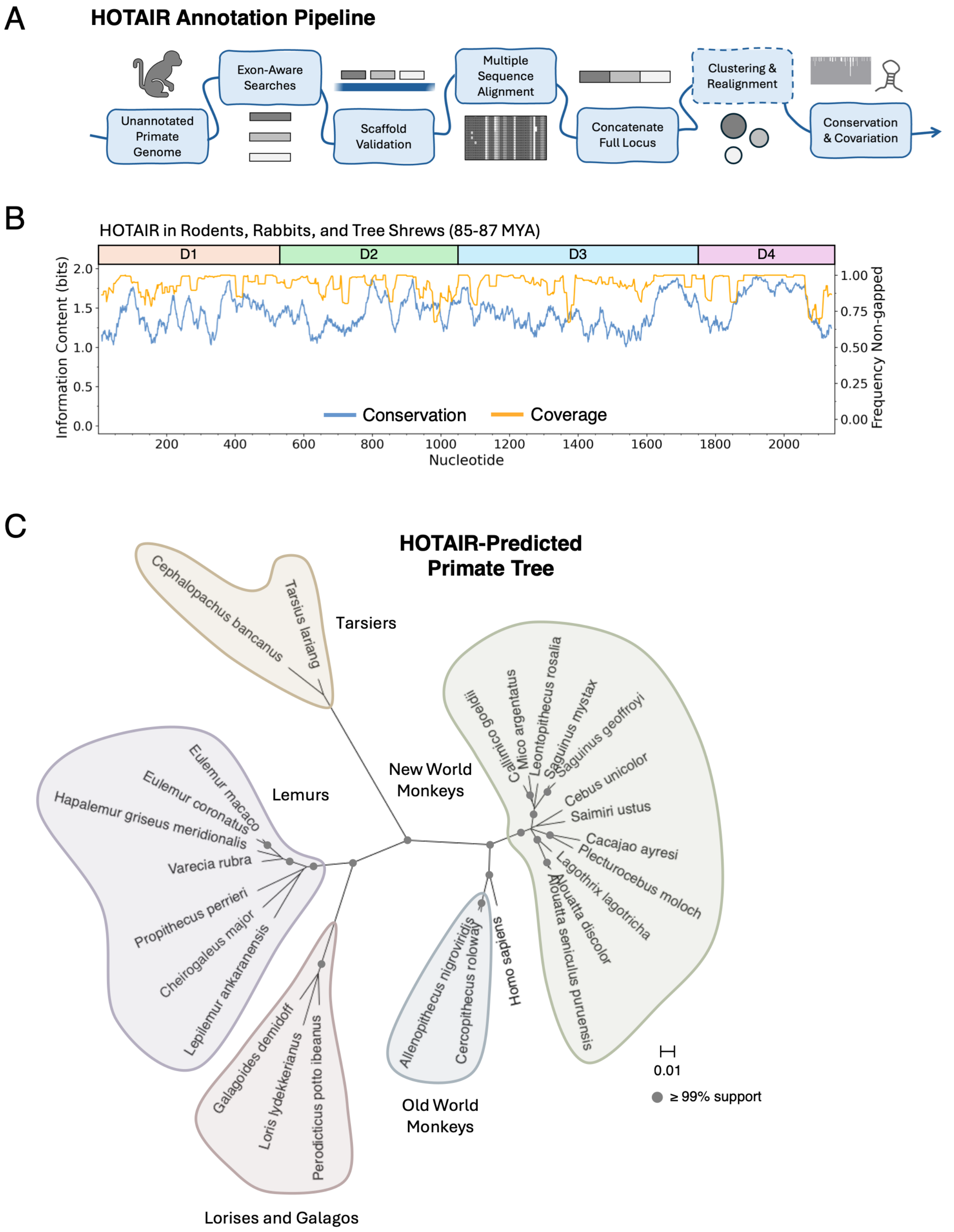


**Supplementary Figure S4.** HOTAIR annotation and conservation in mammals and primates.

(A) Pipeline for annotating the HOTAIR locus in primate genomes, building multiple sequence alignments, and analyzing conservation and covariation. (B) Rolling average of information content and non-gapped frequency across HOTAIR from a multiple sequence alignment of rodents, rabbits, and tree shrews. Domains 1-4 for the *in cellulo* structure are designated as D1-D4. (C) Phylogenetic tree of primate species calculated from the 95% clustered alignment, where 0.01 = 1% sequence divergence. Tree was built using Maximum Likelihood method and GTR+Γ substitution model in UGENE and visualized in MEGA. Filled circles represent branches with ≥ 99% bootstrap support from 1,000 replicates.

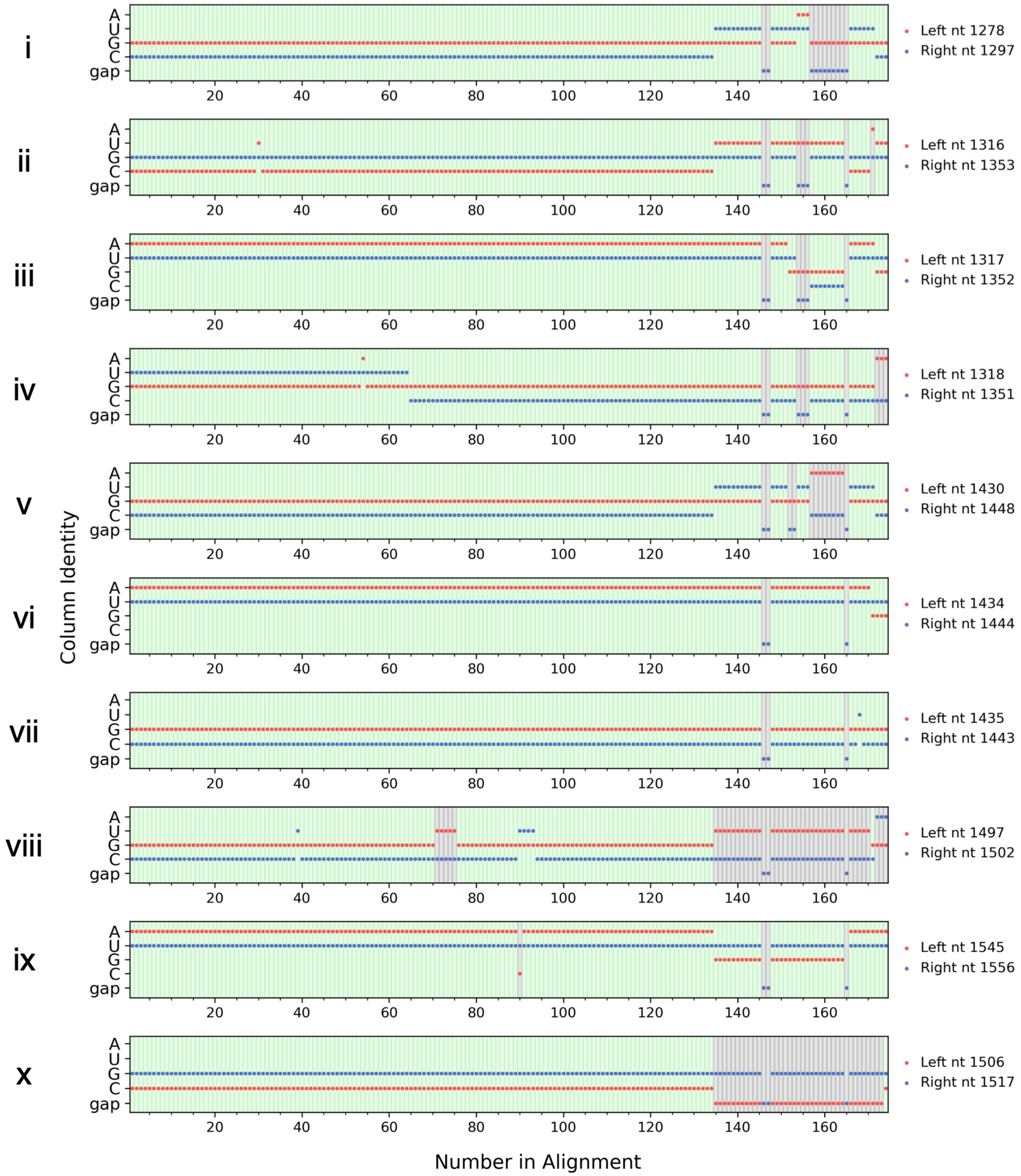


**Supplementary Figure S5.** Compensatory mutations observed in predicted base pairs in HOTAIR Domain 3 across primates. (i-x) All base pairs that were predicted to covary in Domain 3 and were present in the *in cellulo* secondary structure are shown. For each base pair, the column identity (A,

U, G, C, or gap) corresponding to the left nt (red) and right nt (blue) was plotted for each sequence in the pre-clustered alignment of HOTAIR, which contains 174 sequences. Compatible base pairs are highlighted in green, while incompatible base pairs are highlighted in gray. All base pairs exhibit compensatory mutations consistent with covariation, except (x) base pair 1506:1517 that was detected due to high conservation, not covariation. The nt numbers correspond to human HOTAIR.

#### Supplementary Tables

**Supplementary Table S1.** Sequencing quality control metrics of samples from SHAPE-MaP and MRT-ModSeq.

| | Alignment Rate (2A3/DMSO, %)[‡] | Read Depth (%)[§] | Read Depth (nts)[§] | Positive Mutation Rate (%) | Positive Mutation Rate (nts) | High Background Mutation Rate (%) | High Background Mutation Rate (nts) | Highly Reactive (%) | Highly Reactive (nts) |
|---|---|---|---|---|---|---|---|---|---|
| **18S rRNA, *in cellulo* SHAPE-MaP** | | | | | | | | | |
| Rep 1[*] | 94.61/95.32 | 100 | 1231/1231 | 60.4 | 743/1231 | 0.2 | 2/1231 | 9.3 | 114/1231 |
| Rep 2[†] | 94.98/94.48 | 100 | 1231/1231 | 66 | 812/1231 | 0.2 | 2/1231 | 8.6 | 106/1231 |
| **HOTAIR, *in cellulo* SHAPE-MaP** | | | | | | | | | |
| Rep1[*] | 89.45/74.85 | 98.8 | 2049/2074 | 84 | 1721/2049 | 0.1 | 3/2049 | 0.5 | 11/2049 |
| Rep 2[†] | 88.68/81.01 | 98.9 | 2052/2074 | 76.2 | 1563/2052 | 0.1 | 3/2052 | 0.7 | 15/2052 |
| **HOTAIR, *in vitro* SHAPE-MaP**[‖] | | | | | | | | | |
| Rep 1 | 96.63/97.51 | 99.3 | 2061/2075 | 88.5 | 1824/2061 | 0.1 | 3/2061 | 50.3 | 1037/2061 |
| Rep 2 | 95.96/97.27 | 99.5 | 2064/2075 | 88.6 | 1828/2064 | 0.1 | 3/2064 | 48.7 | 1006/2064 |
| **HOTAIR, *in cellulo* MRT-ModSeq**[‖] | | | | | | | | | |
| Rep 1 | 75.88/59.92 | 85.7 | 1778/2074 | 98.9 | 1759/1778 | 0.4 | 8/1778 | 1.6 | 29/1778 |
| Rep 2 | 92.32/94.18 | 98.0 | 2032/2074 | 93.8 | 1907/2032 | 0.4 | 9/2032 | 1.5 | 31/2032 |
| Rep 3 | 82.29/82.51 | 98.2 | 2036/2074 | 97.9 | 1994/2036 | 0.4 | 9/2036 | 5.9 | 121/2036 |

[*,†] Samples marked with the same symbol were derived from the same probed total cellular RNA.

[‡] Primer binding sites were excluded from the calculation of alignment rate and all subsequent metrics. Metrics are the output of ShapeMapper 2 (Busan and Weeks 2018).

[§] The minimum effective read depth was set to the default of 5000 reads.

[‖] Quality control metrics are reported for the complete transcript, which was covered by two overlapping amplicons.

**Supplementary Table S2.** Domain boundaries for *in cellulo*, *in vitro*, and *in silico* HOTAIR secondary structure models.

| Sample | Domain 1 (nt) | Domain 2 (nt) | Domain 3 (nt) | Domain 4 (nt) |
|---|---|---|---|---|
| Replicate 1, *in cellulo* | 1-478 | 479-1055 | 1056-1416 | 1417-2148 |
| Replicate 2, *in cellulo* | 1-525 | 526-1123 | 1124-1795 | 1796-2148 |
| Average, *in cellulo* | 1-530 | 531-1046 | 1047-1747 | 1748-2148 |
| Average, *in vitro* | 1-530 | 531-1047 | 1048-1627 | 1628-2148 |
| Prior structure, *in vitro*[*] | 1-530 | 531-1040 | 1041-1513 | 1514-2148 |
| *in silico* | 1-523 | 524-1122 | 1123-1780 | 1781-2148 |

[*] These domain boundaries were reported for the prior *in vitro* structure (Somarowthu et al. 2015).

**Supplementary Table S3.** Conservation of HOTAIR between human and different taxa.

| Taxon | Divergence Time from Human (MYA) | Number of Species | Median Identity to Human (%) | Median Coverage to Human (%) |
|---|---|---|---|---|
| Eastern Gorilla | 9 | 1 | 98.74 | 99.95 |
| Lesser Apes | 19 | 8 | 94.81 | 98.88 |
| Old World Monkeys | 29 | 54 | 93.30 | 99.53 |
| New World Monkeys | 43 | 70 | 76.50 | 99.12 |
| Tarsiers | 69 | 3 | 76.48 | 96.42 |
| Lemurs, Lorises, and Galagos | 74 | 37 | 78.28 | 98.60 |
| Rodents, Rabbits, and Tree Shrews | 85-87 | 14 | 50.86 | 93.63 |

Percent identity and coverage are reported as the median of pairwise comparisons between each species and human. Values for primates were based on the multiple sequence alignment generated in this study with 174 primates, including human. Values for non-primate Euarchontoglires (rodents, rabbits, and tree shrews) were based on the UCSC 100 Vertebrates MultiZ alignment. Percent identity and coverage were calculated with the NCBI MSA Viewer.

**Supplementary Table S4.** Covarying base pairs detected in Domain 3 from primate alignments of HOTAIR.

| **Left Position (nt)** | **Right Position (nt)** | **E-value[†]** | **FL 99% w500** | **FL 98% w500** | **FL 95% w500** | **D3 95% w500** | **D3 95% w300** | **D3 95% w300 GTp[‡]** |
|---|---|---|---|---|---|---|---|---|
| [*]G1278 | C1297 | 0.037 | | | | | ✓ | |
| [*]C1316 | G1353 | 0.033 | | ✓ | | ✓ | ✓ | |
| [*]A1317 | U1352 | 0.01 | ✓ | ✓ | ✓ | ✓ | ✓ | |
| [*]G1318 | U1351 | 0.023 | ✓ | ✓ | | | | |
| U1397 | A1432 | 0.029 | | | | | ✓ | |
| [*]G1430 | C1448 | 0.033 | | | | ✓ | ✓ | |
| [*]A1434 | U1444 | 0.011 | | ✓ | ✓ | ✓ | ✓ | |
| [*]G1435 | C1443 | 0.037 | | | | ✓ | ✓ | |
| [*]G1497 | C1502 | 0.001 | ✓ | ✓ | ✓ | ✓ | ✓ | ✓ |
| [*]C1506 | G1517 | 0.016 | ✓ | ✓ | ✓ | ✓ | ✓ | |
| G1516 | C1544 | 0.03 | | | | | ✓ | |
| G1529 | C1598 | 0.013 | | | | | ✓ | |
| C1544 | G1578 | 0.042 | | | | | | ✓ |
| [*]A1545 | U1556 | 0.009 | ✓ | ✓ | ✓ | ✓ | ✓ | |

[*] This base pair is supported by the *in cellulo* secondary structure. Although pair C1506:G1517 was detected in the analysis, it is not covarying based on **Supplementary Figure S5**.

[†] The E-value is from the window with the minimum E-value, rounded to 3 significant figures.

‡ This alignment was run with the GTp statistic, whereas all others were run with the RAFSp statistic in R-scape.

A/U/G/C, nucleotide in human HOTAIR; FL, full-length alignment; D3, domain 3 alignment; 99/98/95%, clustering threshold; w500, window size of 500 nt; w300, window size of 300 nt; ✓, base pair was predicted in this alignment.
